# Demographic history and genomics of local adaptation in blue tit populations

**DOI:** 10.1101/864975

**Authors:** Perrier Charles, Rougemont Quentin, Charmantier Anne

## Abstract

Understanding the genomic processes underlying local adaptation is a central aim of modern evolutionary biology. This task requires identifying footprints of local selection but also estimating spatio-temporal variation in population demography and variation in recombination rate and diversity along the genome. Here, we investigated these parameters in blue tit populations inhabiting deciduous *versus* evergreen forests and insular *versus* mainland areas, in the context of a previously described strong phenotypic differentiation. Neighboring population pairs of deciduous and evergreen habitats were weakly genetically differentiated (*F*_ST_ = 0.004 on average), nevertheless with a statistically significant effect of habitat type on the overall genetic structure. This low differentiation was consistent with the strong and long-lasting gene flow between populations, inferred by demographic modeling. In turn, insular and mainland populations were moderately differentiated (*F*_ST_ = 0.08 on average), in line with the inference of moderate ancestral migrations, followed by isolation since the end of the last glaciation. Effective population sizes were overall large, yet smaller on the island than on the mainland. Weak and non-parallel footprints of divergent selection between deciduous and evergreen populations were consistent with their high connectivity and the probable polygenic nature of local adaptation in these habitats. In turn, stronger footprints of divergent selection were identified between long isolated insular versus mainland birds, and were more often found in regions of low recombination as expected from theory. Lastly, we identified a genomic inversion on the mainland, spanning 2.8Mb. These results provide insights into the demographic history and genetic architecture of local adaptation in blue tit populations at multiple geographic scales.

## Introduction

Local adaptation in heterogeneous environments imposing divergent selection on counterparts of a same species has fascinated scientists for decades (Kawecki & Ebert 2004; Blanquart *et al*. 2013). However, it is only with the recent advance in sequencing technologies that numerous recent empirical studies started discovering the genomic features and processes underlying local adaptation in heterogeneous environments (Savolainen *et al*. 2013; Tiffin & Ross-Ibarra 2014; Manel *et al*. 2016; Hoban *et al*. 2016). Especially, many studies used so-called *F*_ST_ genome scans between populations (Excoffier *et al*. 2009; Narum & Hess 2011; de Villemereuil *et al*. 2014) to detect genes under selection, and potentially implicated in local adaptations (Hohenlohe *et al*. 2010; Lamichhaney *et al*. 2016; Reid *et al*. 2016; Perrier *et al*. 2017). Nonetheless, genome scans need improvements for example for studying polygenic variation (Rockman 2012; Wellenreuther & Hansson 2016). Genome scans also need to be interpreted in the light of recombination variation along the genome (Stapley *et al*. 2017, Booker *et al*. 2020) since it shapes the potential extent of diversity and of differentiation in the different regions of the genome (Cutter & Payseur 2013; Tine *et al*. 2014; Burri *et al*. 2015; Rougemont *et al*. 2019). Genome scans results need to be interpreted in the light of the demographic history of populations since i) spatio-temporal variation in gene flow and of effective population size affect the possibility of local adaptations (Lenormand 2002) and the level of false positive due to genetic drift, and ii) regions implicated in reproductive isolation accumulated during allopatry showing increased differentiation upon secondary contacts can be misinterpreted as genuine footprints of recent divergent selection (Bierne *et al*. 2011; 2013). Lastly, structural variants such as inversions also require special attention, since they protect adaptive gene sets from recombination (Kirkpatrick 2006; 2010; Stapley *et al*. 2017; Wellenreuther *et al*. 2019) and hence enable their persistence and rapid redeployment in heterogeneous environments despite high gene flow (Lowry & Willis 2010; Sinclair-Waters *et al*. 2017; Todesco *et al*. 2019).

Here, we aimed at investigating various demographic and genomic aspects of the adaptive divergence of a well-studied passerine bird, the blue tit (*Cyanistes caeruleus*). Populations of small insectivorous passerines have long been used to study local adaptation, in both quantitative genetics and population genetics frameworks (Carbonell *et al*. 2003; Broggi et al. 2005; Laaksonen et al. 2015). In particular, several blue tit populations breeding in a heterogeneous habitat in Southern France (Figure 1A) offer an ideal context to study local adaptation. Four of them (two deciduous and two evergreen, circled in black in Figure 1A) have been subject to a long term project spanning more than 40 years (Blondel et al. 2006; Charmantier et al. 2016). These populations show marked quantitative phenotypic differences (Figure 1B&C), notably in morphological (e.g. tarsus length and body mass), life-history (lay date and clutch size) and behavioural traits (e.g. song characteristics and handling aggression) (Charmantier et al 2016). These phenotypic differences were found at two spatial scales. First, birds breeding in deciduous forest habitats are taller, more aggressive, and lay larger and earlier broods than birds in evergreen forests (see table 1 in Charmantier et al 2016). Strikingly, neighbouring populations in deciduous and evergreen habitats are weakly genetically differentiated (Porlier et al. 2012b; Szulkin et al. 2016; Dubuc-Messier *et al*. 2018) despite the short spatial scale, which questioned the mechanisms of persistence of the observed phenotypic differentiation against presumably large gene flow. Second, insular blue tits from Corsica, that might have been isolated since the sea level rise after the last glacial maximum (the sea level raised of 120 m from 17,000 to 5,000 years ago (Lambeck & Bard 2000; Jouet et al. 2006)), are smaller and more colourful than their mainland counterparts (again, see table 1 in Charmantier *et al* 2016) and are listed as two subspecies (*Cyanistes caeruleus caeruleus* on mainland Europe and *Cyanistes caeruleus ogliastrae* mainly in Corsica and Sardinia). Overall, traits displaying these strong phenotypic differences had heritability ranging from 0.20-0.43 (eg for lay date, (Caro et al. 2009)) to 0.42-0.60 (eg for tarsus length, (Teplitsky et al. 2014; Delahaie et al. 2017; Perrier et al. 2018)), are classically related with fitness, and hence could be involved in a local adaptation process. In particular, the heritability of lay date and the breeding time gap between populations could result in reproductive isolation by breeding time, limiting gene flow and favouring local adaptation. The studied traits were typically quantitative (Charmantier et al. 2016), hence probably controlled by a polygenic architecture (Perrier et al. 2018) composed of many loci with small individual effects, as found in similar traits for other passerine birds (Santure et al. 2013; Bosse et al. 2017; Hansson et al. 2018; Lundregan et al. 2018). Overall, given their phenotypic, demographic and genetic characteristics, these blue tit populations provide an ideal framework to study the genomic architecture of polygenic adaptation in heterogeneous environments.

**Table 1.** Study populations’ names, abbreviations, habitat types, locations, approximate forest area (ha), altitude (m), and sample sizes.

| Name | Abbreviation | Region | Site | Type of<br>oaks | Latitude | Longitude | Forest<br>area | Altitude | N-GeneralDataset | N-FilteredDataset |
| --- | --- | --- | --- | --- | --- | --- | --- | --- | --- | --- |
| D-Rouviere | A1D | Mainland | Rouviere | Deciduous | 43.664000 | 3.668000 | 465 | 248 | 203 | 60 |
| D-Quissac | A2D | Mainland | Quissac | Deciduous | 43.891036 | 3.996931 | 23 | 104 | 35 | 27 |
| E-Quissac | A2E | Mainland | Quissac | Evergreen | 43.926673 | 3.989759 | 415 | 116 | 31 | 24 |
| E-Pirio | B3E | Corsica | Pirio | Evergreen | 42.376000 | 8.750000 | 400 | 145 | 180 | 60 |
| D-Muro | B4D | Corsica | Muro | Deciduous | 42.551000 | 8.923000 | 188 | 283 | 215 | 60 |
| E-Muro | B4E | Corsica | Muro | Evergreen | 42.589000 | 8.967000 | 80 | 99 | 116 | 60 |
| D-Zilia | B5D | Corsica | Zilia | Deciduous | 42.523685 | 8.873689 | 13 | 102 | 42 | 38 |
| E-Zilia | B5E | Corsica | Zilia | Evergreen | 42.520633 | 8.884057 | 17 | 267 | 38 | 38 |
| D-OlmiCappella | B6D | Corsica | OlmiCappella | Deciduous | 42.540078 | 8.993915 | 20 | 1027 | 44 | 44 |
| E-OlmiCappella | B6E | Corsica | OlmiCappella | Evergreen | 42.529414 | 9.021652 | 25 | 881 | 43 | 43 |
| Total/average |  |  |  |  |  |  |  |  | 947 | 454 |

**Figure 1.**
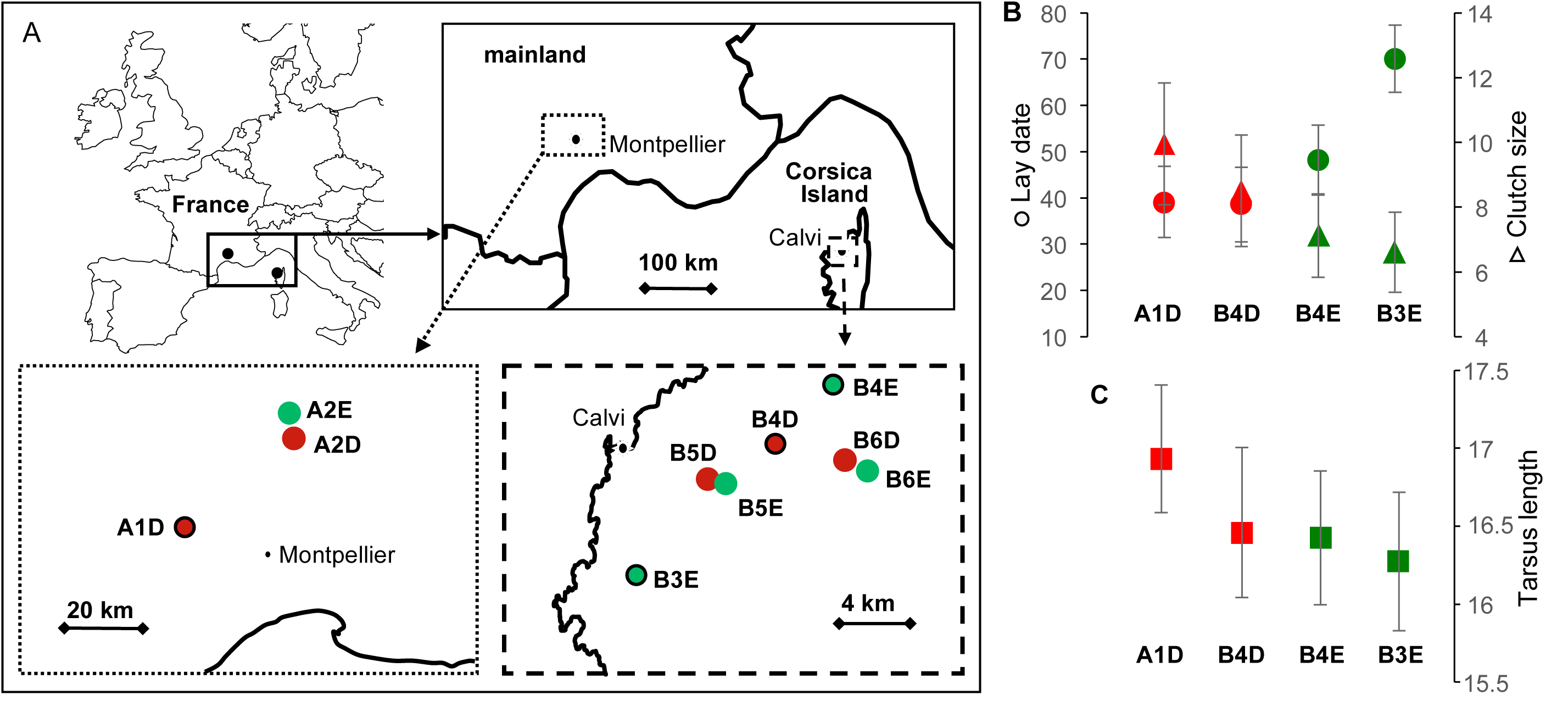
Map of the sampling locations of blue tit populations on the mainland and in Corsica (A) and phenotypic differences (B). In red, habitats dominated by deciduous oaks and in green, habitats dominated by evergreen oaks. Sites with long-term monitoring are circled in black on the map (A1D is D-Rouvière, B3E is D-Pirio, D4D is D-Muro and B4E is E-Muro). On figures B & C are presented illustrative phenotypic differences (with standard deviations) between the four main populations with long term monitoring. Are shown laying date (days), clutch size and male tarsus length (mm) (data from Table 1 in Charmantier et al 2016 Evol Appl, see this reference for detailed information for these measures and for differences in many other traits).

We investigated genome wide patterns of genetic diversity and differentiation and the demographic history between several populations of blue tit from Southern France, in heterogeneous forest habitats (deciduous versus evergreen) and in insular (Corsica island) and mainland areas (Mainland France) in order to better understand the determinants of their local adaptation. The analysis was based on Resequencing of birds sampled in four sites studied in the context of a long-term project (Blondel et al. 2006; Charmantier et al. 2016) together with three additional pairs of deciduous and evergreen forests in order to test for parallel evolution (see Figure 1A). First, we investigated variation in genetic diversity and differentiation in order to verify that habitat type and geographic distance explained a significant proportion of the genetic structure between populations (Szulkin et al. 2016). Second, we investigated the historical and contemporary demography of each deciduous and evergreen population pair in order to better understand the origin of their differentiation, and the demography between the populations on mainland France and on the Corsica Island in order to estimate their split time and their subsequent connectivity and effective population size. Third, we screened the genome for *F*_ST_ outlier loci and inversions potentially implicated in adaptation in the two habitat types and between Corsica and Mainland France. Rather than identifying the genes under selection, we wanted to test whether the adaptation had a polygenic or oligogenic architecture and whether outlier loci would be more frequent in regions with low recombination rate. We discuss our results in the light of the recent literature in population genomics, to decipher the role of genomic and demographic variations on the genetic and phenotypic divergence between blue tits in deciduous *versus* evergreen habitats and in mainland versus insular contexts.

## Methods

### Sites and sampling

947 blue tits (*Cyanistes caeruleus)* were captured in ten locations (Figure 1A, Table 1). Three locations are located in the South of mainland France (codes staring with “A”: A1D, A2D, A2E) and seven in the Corsica Island (staring with “B”: B3E, B4D, B4E, B5D, B5E, B6D, B6E). The numbers from 1 to 6 indicated the sampling area and hence the population pairs. Half of these sites were dominated by the deciduous downy oak *Quercus pubescens* (indicated by “D” in the code) and the other half are dominated by the evergreen holm oak *Q. ilex* (indicated by “E”). Four of these locations (A1D, B3E, B4D, B4E) are monitored as part of a long-term research program started in 1976 and have been described in previous studies (Blondel et al. 2006; Charmantier et al. 2016). The 6 other sites (A2D, A2E, B5D, B5E, B6D & B6E) were sampled in order to obtain further replicates of neighboring deciduous and evergreen populations. Capture and handling of the birds was conducted under permits provided by the *Centre de Recherches sur la Biologie des Populations d’Oiseaux* (CRBPO) and by the *Direction Départementale des Services Vétérinaires* (DDSV). Birds were captured during the reproductive period, from early April to late June, on their nesting territory. Birds were captured either in nest boxes during the feeding of their young (in the four sites studied on the long term), or using nets in the vicinity of their nest (in the other sites). Birds were banded with a unique metal ring provided by the CRBPO. Five to 20μl of blood were sampled from a neck or a wing vein from breeding blue tits. Blood was stored at 4°C in Queen’s buffer (Seutin *et al*. 1991).

### Molecular biology and sequencing

DNA extractions were achieved using Qiagen DNeasy Blood & Tissue kits and were randomized across sites. DNA was quantified using first a NanoDrop ND8000 spectrophotometer and then a Qubit 2.0 fluorometer with the DNA HS assay kit (Life Technologies). DNA quality was examined on agarose gels. Library preparation using RAD-seq (restriction-site-associated DNA sequencing; (Baird et al. 2008)) with the enzyme *SbfI* was done by Montpellier GenomiX (MGX) platform (CNRS, Montpellier). Each individual was identified using a unique 6 nucleotides tag, individuals were multiplexed in equimolar proportions by libraries of 36 individuals, and each library was sequenced on a lane of an Illumina HiSeq 2000. Single-end sequencing was used to produce 100bp reads. This design was used to obtain an average read depth of 50x. The DNA of three individuals was extracted twice and processed in different libraries to test for reliability of the genotyping process.

### Bioinformatics and data filtering

Raw sequences were inspected with *FastQC* (Andrews 2010) for quality controls. Potential fragments of Illumina adapters were trimmed with *Cutadapt* (Martin 2011), allowing for a 10% mismatch in the adapter sequence. Reads were filtered for overall quality, demultiplexed and trimmed to 85bp using *process_radtags*, from the *Stacks software pipeline* 1.39 (Catchen et al. 2013), allowing for one mismatch in the barcode sequence. Sequencing RAD-tags resulted in a median value of 5,449,564 reads per individual. *BWA-MEM* 0.7.13 (Li & Durbin 2009) was used to map individual sequences against the reference genome of the great tit *Parus major* (Laine et al. 2016) and to produce *sam* files using default options. Indeed, although great tit and blue tit diverged about 7 to 14My ago (Päckert et al 2007), the use of a reference genome over a de-novo approach is very often highly recommended (See e.g. Rochette et Catchen 2017). It is also interesting to note that synteny is typically highly conserved in birds (Backström et al 2008), suggesting that aligning reads on a divergent species would still allow analyses based on SNP position along the genome. On average, 93% of the raw reads were mapped against the genome (Supplementary table 1). *Samtools* 0.1.19 (Li et al. 2009) was used to build and sort *bam* files. We used pstacks to treat *bam* files, align the reads as assembled loci and call SNPs in each locus. We used a minimum depth of coverage (m) of 5, the SNP model, and alpha = 0.05 (chi square significance level required to call a heterozygote or homozygote). *cstacks* was used to build the catalogue of loci, allowing 3 mismatches between sample loci when building the catalog. *sstacks* was used to match loci against the catalog. Lastly, *populations* program in Stacks was used to genotype individuals. In this program, relatively permissive filters were applied, to retain SNP genotyped in at least 50% of individuals (all individuals from all sites grouped) and with a minimum read depth of 4. 350,941 SNP and 947 individuals were obtained in this “GeneralDataset”.

We hence applied additional filters to this GeneralDataset using the programs *VCFtools* (Danecek et al. 2011) and *Plink* (Purcell et al. 2007). We filtered for a minimum average read depth of 10 and a maximum average read depth of 100 (corresponding approximately to the 5%-95% distribution of read depth). SNP genotyped in less than 80% of the individuals were removed. We removed SNPs with observed heterozygosity ≥ 0.65 among individuals on Corsica or on mainland to reduce the potential occurrence of stacked paralogues. Individuals below 85% genotyping rate were removed. Identity-By-State (IBS) was estimated for three replicated individuals (from the DNA extraction to SNP-calling) in order to investigate reliability of the entire genotyping process. The IBS measured between replicates were high, ranging from 0.9989 to 0.9999, indicating very low genotyping error rate. These replicates were then removed from the dataset. Using the R packages *gdsfmt* and *SNPRelate* (Zheng et al. 2012), we measured the realized genomic relatedness (GRM) between individuals in order to remove highly related individuals (*i*.*e*. full-sibs and parent-offspring). For each pair of individuals with relatedness ≥ 0.35, we removed one individual. This procedure was applied in order to limit biases due to highly related individuals. Indeed, since we sampled breeding birds at their nests for the four sites studied on the long-term and that they tend to disperse relatively close to their nests, we expected a higher percentage of related individuals than there actually is in the population. We then limited the number of individuals to 60, chosen at random, in populations (A1D, B3E, B4D & B4E) in which a large number of individuals were genotyped. This limitation was achieved in order to limit differences in analyses precision between populations due to unequal sample sizes (all individuals will be used in another ongoing study of the genomic architecture of quantitative adaptive traits). We hence removed potential monomorphic SNPs and created the dataset “FilteredDataset”. 454 individuals (Table 1) and 144,773 SNPs were kept in this final dataset used for all the genomic analyses excepted analyses of demographic history using ABC. The median genotyping rate across all SNPs for these individuals was 0.981. The median read depth across genotypes (SNPs x Individual) was 49.2x. The number of SNPs per chromosome ranged from 143 (LGE22) to 15,738 (chromosome 2). Further filtering (e.g. MAF) was often operated, depending on the analyses, and thereafter mentioned if it was the case. To perform analyses of demographic history using ABC analyses, we produced a haplotype VCF file for these 454 aforementioned individuals with the populations module of Stacks and we filtered it as explained in the supplementary note 1.

### Analysis of genetic structure and effects of environmental variables

We used PCAs with the function *snpgdsPCA* from *snprelate* to depict genetic structure between individuals. PCAs were run for the entire dataset, for mainland and Corsican populations separately and for each pairs of deciduous and evergreen populations. We inferred admixture proportions for each individual using *Admixture* 1.23 (Alexander et al. 2009) with K-values ranging from 1 to 12 and 1,000 bootstraps. The different clustering solutions were inferred by plotting cross-validations errors and by plotting individual admixture proportion. The effect of environmental variables, forest phenology (E versus D), geographic distance (latitude & longitude) and altitude, on genomic differentiation was measured using a redundancy analysis (RDA) (Legendre & Fortin 2010; Forester et al. 2015) as implemented in the R package *Vegan* (Oksanen et al. 2007). We investigated the proportion of genetic variability explained by a constraining covariance matrix consisting of phenology, latitude, longitude, and altitude for each individual. We tested the global significance of the model using 1,000 permutations. We ran marginal effects permutations to address the significance of each variable. Then we focused on the effect of phenology alone, using partial RDA to take into account the effect of latitude, longitude and altitude. Significance was tested running 1,000 permutations. For the PCA, the admixture analysis and the RDA, we selected from the FilteredDataset the SNPs with more than 95% genotyping rate, MAF > 0.05, retaining one SNP per locus and we remove SNP in linkage disequilibrium using the *Plink* command “indep 50 5 2”. Genome wide differentiation between each sampling location was measured with Weir and Cockerham’s *F*_ST_ estimator (Weir & Cockerham 1984) implemented in *StAMPP* (Pembleton et al. 2013). *F*_ST_ was estimated for all the SNPs, the ones on the autosomes and the ones on the sex chromosome Z separately. Significance was assessed using 1,000 bootstraps replicates.

### Analysis of demographic history

Alternative models of divergence history including the effects of selection at linked sites affecting *Ne* and of differential introgression (*m*) were compared using an ABC framework modified from Roux et al. (2016). Linked selection either under the form of hitchhiking of neutral loci linked to a selective sweep (Maynard Haigh & Smith, 1974) or under the form of background selection (Charlesworth et al. 1993), have strong effects in regions of low recombination and have been shown to influence model choice and parameter estimates (Ewing & Jensen, 2015; Schrider et al. 2016). The same is true when populations accumulate reproductive incompatibilities during the divergence process: the resulting barrier to gene flow reduces the effective migration rate along the genome (Barton & Bengtsson, 1986) and not accounting for it can affect demographic model choice and parameter estimates (Roux et al. 2014; Souza et al. 2013). Moreover, including selected loci in demographic inferences can reveal the deeper origins of population divergence (Bierne et al. 2013). Six scenarios were compared for the 4 pairs of deciduous and evergreen populations in order to test whether the divergence between different habitats was not due to a divergence in different historical refugia but to a contemporary ecological divergence. We included a model of panmixia (PAN), a model of equilibrium corresponding to the island model with two populations (EQ), a model of isolation with migration (IM), a model of secondary contact (SC), a model of divergence with migration during the first generations, *i*.*e*. ancestral migration (AM), and a model of strict isolation (SI). The prior and details of the simulation pipeline are fully described in the supplementary note 1. The PAN model assumes that the two focal populations descent from a single panmictic population characterized by its effective size (Ne). The EQ model (equivalent to the island model) assumes that the population is subdivided into two discrete populations of sizes *N*_pop1_ and *N*_pop2_ that are connected by continuous gene flow at a constant rate each generation. In this model, the divergence time is not a parameter. The IM, SI, SC, and AM models all assume that an ancestral population of size *N*_ANC_ splits at *T*_*split*_ into two daughter populations of sizes *N*_pop1_ and *N*_pop2._ Under SI no subsequent gene flow occurs. Under AM model, gene flow occurs from *T*_*split*_ to *T*_am_ and is followed by a period without gene flow. Under IM, gene flow is continuous after *T*_*split*_. Under, the SC model, *T*_*split*_ is followed by a period of strict isolation, after which a secondary contact starts *T*_sc_ generations. The EQ, IM, AM and SC models included migration as *M* = 4 *N*_*0*_.*m*, with M_1 ←2_ being the number of migrants from population 2 to population 1 and M_2 ←1_ being the reverse. The effect of linked selection and barriers to gene flow were accounted for, by modeling heterogeneous population size (Ne) and heterogeneous migration (m) respectively. Such heterogeneity was modeled using beta distributions as hyper-prior on each of these two parameters. These resulted in four alternative versions for models with gene flow EQ, AM, IM, SC (NhomoMhomo, NhomoMhetero, NheteroMhomo, NheteroMhetero) and two versions for PAN and SI (Nhomo and Nhetero). We used a modified ABC (Csilléry et al. 2010) pipeline from (Rougemont & Bernatchez 2018) to perform model selection, compute robustness and to estimate posterior probabilities of parameters.

We also investigated the historical demography of the populations from Corsica as compared to the ones from the mainland. Gene flow has probably been impossible since the last de-glaciation and sea-level rise. Therefore, we compared models of ancient migration (AM) and Strict Isolation (SI). The pipeline described in the above section, integrating linked selection and barriers to gene flow, was run between the A1D samples (chosen on the mainland because it had the largest sample size) and B4D (chosen in Corsica because it had both a large sample size and was from the same habitat as A1D). We used the same ABC pipeline for the model selection procedure and parameter estimation as described in the above section. Finally, we attempted to convert demographic parameter into biological units assuming a mutation rate of 1e-8 mutations/bp/generations.

### Genomic diversity

Genome wide genetic diversity was inferred for each sampling location in the dataset by measuring observed heterozygosity (Ho), proportion of polymorphic loci and MAF spectrums. For each population and chromosome, and subsequently on average for the entire genome, Linkage Disequilibrium (LD) decay was measured with *Plink* and smoothed in R. To contrast with long term Ne estimates from coalescent simulation in our ABC modeling (see “Analysis of demographic history”), we also inferred recent Ne for each population using *SNeP* V1.1 (Barbato et al. 2015), which uses LD data, with a MAF ≥ 0.05 filter per population. We investigated the nature of SNP variation, *i*.*e*. synonymous or non-synonymous, blasting all rad sequences on the reference genome and the transcriptome of the great tit (ftp://ftp.ncbi.nih.gov/genomes/Parus_major/; Santure et al 2011) using *blastx* (McGinnis & Madden 2004). We kept hits with at least 90% similarity and a minimum amino acid sequence length alignment of 25. We kept only SNPs for which both the alternative and reference allele yields the same score. Finally, we tested for differences in the distribution of run of homozygosity (ROH) between the mainland and Corsica that may have resulted from smaller Ne and larger inbreeding in Corsica versus the mainland. We used plink 1.9 to estimate the length and number of ROH. We required a window of 500 kb to be homozyguous in order to be considered as a ROH, and with a maximum of 100 SNP using the following parameters: -- homozyg-density 50,--homozyg-gap 100,--homozyg-kb 500,--homozyg-snp 100,-- homozyg-window-het 1,--homozyg-window-snp 100,--homozyg-window-threshold 0.05, and--homozyg-window-missing 20.

### Identification of genomic footprints of selection

We used three methods to search for outlier SNPs potentially under divergent selection between blue tit populations. First, we used *Bayescan* V2.1 (Foll & Gaggiotti 2008) to search for SNPs potentially under divergent selection at two geographic levels: i) between the mainland and Corsican populations and ii) between each local pairs of evergreen (E) and deciduous (D) populations, *i*.*e*. A2D vs A2E, B4D vs B4E, B5D vs B5E, B6D vs B6E. We filtered each of the five datasets for a minimum MAF of 0.05. We used default parameters except for prior odds that were set at 10,000 in order to limit false positives. We investigated the parallelism across pairs of D-E environments. Second, we estimated *F*_ST_ along the genome using either a 200kb sliding average with *VCFtools*, or the function “snpgdsSlidingWindow” from the package SNPrelate to estimate *F*_ST_ in blocks of 50 SNPs moving by 5 SNPs. The second window strategy was used in order to compensate for lower SNP density in regions of low recombination that tend to exaggerate the contribution of individual SNPs in these regions and to dilute the individual SNP contribution in regions of high recombination (see Perrier & Charmantier 2019 for a broader comment on this). Third, we used an RDA as an alternative method to search for SNPs putatively implicated in multilocus adaptation i) between populations in deciduous and evergreen habitats, and ii) between populations in Corsica and on the mainland. As suggested by Forester et al (2018), such a multivariate method may be more suitable than univariate ones to detect weaker footprints of adaptation that are expected in polygenic adaptations in response to complex environmental heterogeneity. Using a similar procedure as described earlier in the methods, we used two RDAs constrained to investigate the effect of phenology (i) or of the geography (ii). We then used a 3 standard deviation cutoff as suggested by Forester et al (2018) to list loci with outlier loading scores on the first RDA axes. We compared the loci found using these different methods. We reported in which genes these outliers were found (the list of genes can be found together with the genome published by Laine et al 2016 on NCBI).

### Variation of genomic differentiation with recombination rate

We investigated variation of F_ST_ with local recombination rate and whether SNP outliers were more often found in regions of low recombination than elsewhere in the genome. We estimated local recombination rates using a coalescent method implemented in *Ldhat* (McVean 2004) using linkage disequilibrium signal. Following these authors’ recommendation, the dataset was split in blocks of 2,000 SNPs with 500 overlapping SNPs. A MAF of 0.05 and a maximum of 5% of missing data were allowed. The local recombination rate, ρ = 4Ne r, was then estimated in each block independently with the Bayesian reversible jump MCMC scheme implemented in interval. We used a block penalty of 5, with 30 millions MCMC iterations sampled every 5,000 iterations. The first 250,000 iterations were discarded. To speed up computations we used the pre-computed two locus likelihood table for n = 190 and assuming theta = 0.001. We estimated ρ in a composite dataset with individuals from every population, habitats and sex. We tested for correlations between SNP *F*_ST_ and recombination rate using linear models, for Corsica-mainland and for deciduous-evergreen differentiation, independently, and we represented the correlation using a LOESS fit. We tested whether outliers found using Bayescan and the RDA method for both Corsica-mainland and deciduous-evergreen differentiation were more often found in regions of low recombination than elsewhere in the genome using chi-square (χ2) tests, and we represented the pattern using histograms.

### Detection of genomic inversions

We searched for potential genomic inversions using a variety of descriptive statistics. First, we searched for genomic regions having a particularly low recombination rate and large long-distance linkage disequilibrium, nevertheless associated to a high density of SNPs and therefore unlikely to correspond to pericentromeric regions but rather to local suppression of recombination that may be due to inversions. Second, we implemented a PCA sliding window analysis in order to identify portions of the genome with individuals carrying an inversion at the homozygous or heterozygous state or individuals exempt from the inversion (Ma & Amos 2012). We first used 10Mb windows sliding by 1Mb and then 1Mb windows sliding by 100kb, in order to use enough SNPs to perform PCAs. We then used *Lostruct* (Li & Ralph 2019), with k=2, sliding by 100 SNPs, to identify particular blocks of linked SNPs explaining an abnormally high proportion of variance between 2 groups of individuals (e.g. inverted and non-inverted).

We detected one putative inversion. We verified, using admixture with K = 2 for analyzing SNPs from the inversion, that putative heterozygous individuals had an admixture ratio close to 1:1 of both putative inverted and non-inverted homozygous clusters. We then inspected variations of *F*_ST_ (per SNP) and π (per 10kb window, using *vcftools*) along the genome between individuals that were inverted homozygous, non-inverted homozygous, and heterozygous for the inversion, looking for potentially reduced diversity and increased differentiation at the inversion. We looked for potential salient variations in read depth in and around the putative inversion. We also tested whether the detected inversion was at Hardy-Weinberg equilibrium (using a common χ2-test) and whether the frequency of the inversion varied geographically and between evergreen and deciduous habitats.

To study the history of the putative inversion identified, we aimed at measuring intra- and inter-specific genetic distance and absolute divergence, at the inversion and for the entire genome, for several blue tits and great tits. To do that, we first generated a new SNP dataset by running stacks with the same pipeline and parameters as explained earlier, with 2 mainland great tits, 2 insular great tits, 4 mainland blue tits homozygous for the inversion, 4 mainland blue tits not carrying the inversion, and 4 Corsican blue tits not carrying the inversion (the inversion was not found in Corsica). Second, we selected the SNPs from the region of the inversion. Third, Dxy was measured between the 5 aforementioned groups of individuals using *PopGenome* (Pfeifer et al 2014). Using this measure of Dxy, we estimated approximately the inversion apparition time using T = Dxy / 2µ, assuming a standard mutation rate of 1e-8 and with the simplifying assumption of no gene flow and no introgression. Lastly, we represented the divergence between these individuals using an UPGMA tree of bitwise distance using the R package *poppr* 1.1.1 (Kamvar et al 2014).

### Gene ontology

We used the R package topGO (Alexa & Rahnenführer, 2009) to investigate the potential gene ontologies (GO) that were statistically enriched for the sets of genes identified among outliers and the inversion, compared to the entire list of genes in which all the SNPs from the entire dataset were found. For the outlier tests, we used GO analyses for each gene list obtained using the different outlier identification methods but also for aggregated lists of all of the gene identified for deciduous-evergreen tests or for Corsica-Mainland tests. We used the GO referenced for the zebra finch, T. Guttata (tguttata_gene_ensembl). We report a Fisher enrichment test p-value and a p-value after applying a Benjamini and Hochberg (BH) correction for multiple testing to control for false discovery rate (FDR).

## Results

### Genetic structure and effects of environmental variables

The admixture analysis suggested the existence of 2 main distinct genetic groups corresponding to mainland and island populations (Figure 2A). Increasing K value contributed to delineate the populations, showing the existence of a weak structure between the 6 main areas. However, the coefficient of variation increased with K. There was very little evidence for genetic structure within each of the four pairs of D- and E-populations. PCAs revealed a clear structure between individuals from the mainland and the island but little structure within each of these 2 groups (Supplementary Figure 1). The full RDA model was significant (p < 0.001, Supplementary table 2, Figure 2B), as well as the effects of each variable tested in the model: phenology (*i*.*e*. E versus D, p = 0.006), latitude (p < 0.001), longitude (p < 0.001) and altitude (p < 0.001). The first axis of the global RDA explained 4.54% of the variance, and was correlated mainly with latitude (- 0.987) and longitude (+0.999). The fourth axis of this global RDA displayed the strongest correlation with phenology (−0.856), and explained 0.27% of the variance. The partial RDA model conditioning for the effects of latitude, longitude and altitude, was globally significant (p < 0.001), as well as the effect of phenology (p <0.001). The axis of the partial RDA explained 0.29% of the variance and was correlated with phenology (0.922).

**Table 2.** Diversity and differentiation across populations. Fixation index between populations (*F*_ST_), effective population size (Ne) estimated with ABC (Ne1) using models for Deciduous-Evergreen population pairs or for Corsica-Mainland (A1D-B4D), Ne estimated with a LD method (Ne2), and observed heterozygosity (Ho). Color gradients indicate differentiation intensity (Darker red for higher differentiation and darker blue for lowest differentiation). Underlined values indicate deciduous and evergreen pairs.

| | $F_{ST}$ | | | | | | | | | | Ne1 | | Ne2 | Ho |
| --- | --- | --- | --- | --- | --- | --- | --- | --- | --- | --- | --- | --- | --- | --- |
|  | A1D | A2D | A2E | B3E | B4D | B4E | B5D | B5E | B6D | B6E | Deciduous-Evergreen | Corsica-Mainland |  |  |
| A1D |  | 0.0045 | 0.0043 | 0.0812 | 0.0830 | 0.0853 | 0.0796 | 0.0804 | 0.0785 | 0.0784 | Na | 397,685 | 355 | 0.171 |
| A2D | 0.0045 |  | 0.0029 | 0.0832 | 0.0849 | 0.0872 | 0.0799 | 0.0810 | 0.0793 | 0.0792 | 400,005 | na | 161 | 0.167 |
| A2E | 0.0043 | 0.0029 |  | 0.0828 | 0.0846 | 0.0871 | 0.0796 | 0.0807 | 0.0790 | 0.0787 | 100,055 | na | 142 | 0.168 |
| B3E | 0.0812 | 0.0832 | 0.0828 |  | 0.0058 | 0.0089 | 0.0067 | 0.0071 | 0.0056 | 0.0058 | Na | Na | 334 | 0.149 |
| B4D | 0.0830 | 0.0849 | 0.0846 | 0.0058 |  | 0.0079 | 0.0063 | 0.0067 | 0.0061 | 0.0062 | 43,070 | 52,990 | 310 | 0.149 |
| B4E | 0.0853 | 0.0872 | 0.0871 | 0.0089 | 0.0079 |  | 0.0093 | 0.0097 | 0.0083 | 0.0085 | 37,100 | na | 271 | 0.148 |
| B5D | 0.0796 | 0.0799 | 0.0796 | 0.0067 | 0.0063 | 0.0093 |  | 0.0025 | 0.0035 | 0.0032 | 46,520 | na | 210 | 0.145 |
| B5E | 0.0804 | 0.0810 | 0.0807 | 0.0071 | 0.0067 | 0.0097 | 0.0025 |  | 0.0036 | 0.0033 | 42,800 | na | 216 | 0.146 |
| B6D | 0.0785 | 0.0793 | 0.0790 | 0.0056 | 0.0061 | 0.0083 | 0.0035 | 0.0035 |  | 0.0006 | 46,885 | na | 268 | 0.146 |
| B6E | 0.0784 | 0.0792 | 0.0787 | 0.0058 | 0.0062 | 0.0085 | 0.0032 | 0.0033 | 0.0006 |  | 162,940 | na | 260 | 0.146 |

**Figure 2.**
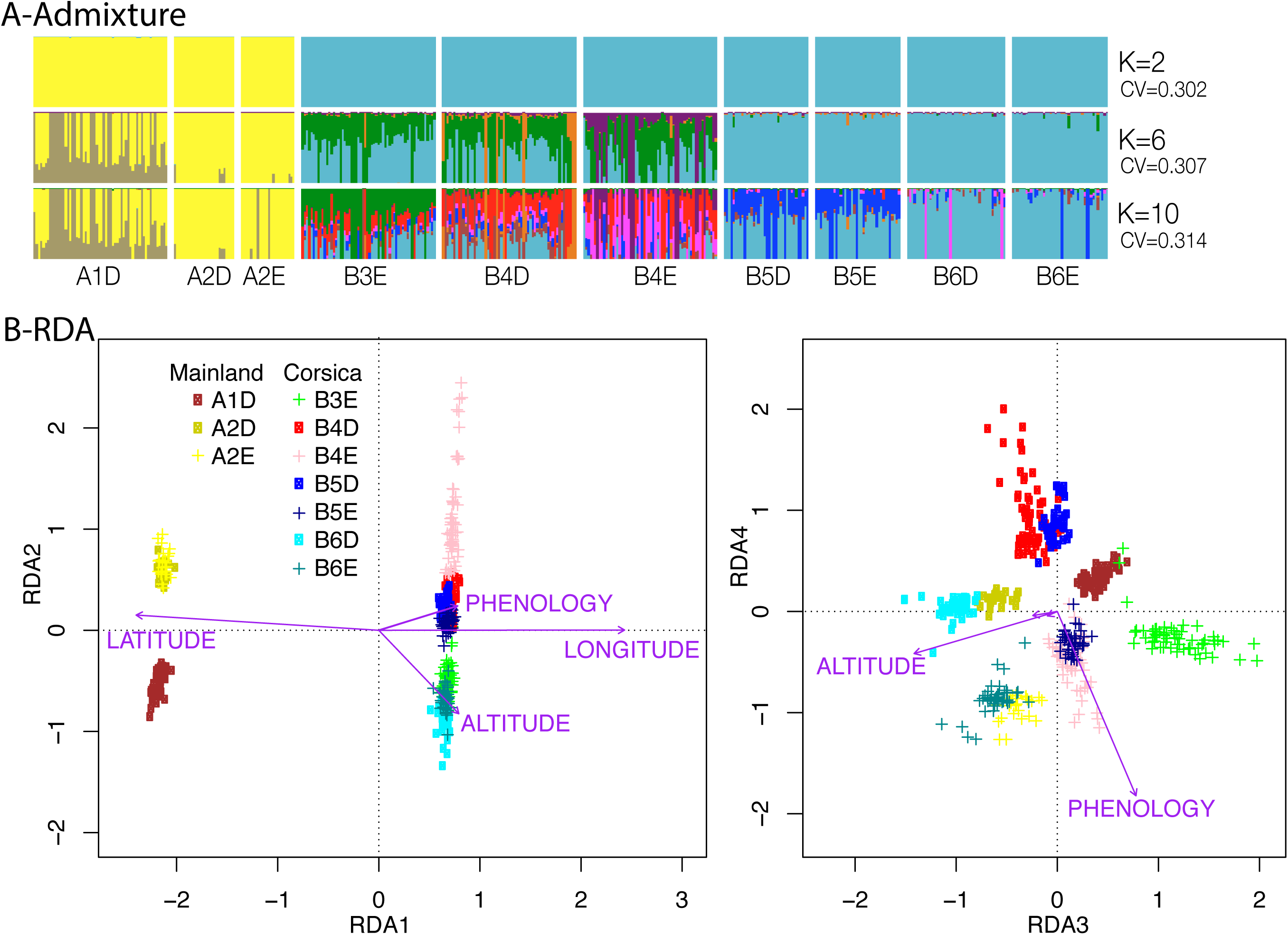
Population structure between populations. **A)** Admixture for K = 2, 6 and 10 (Vertical lines correspond to individual admixture; colors correspond to distinct genetic groups). **B)** RDA showing the influence of latitude, longitude, altitude and phenology on genetic structure.

All pairwise *F*_ST_ comparisons were significantly different from zero (Table 2). Average *F*_ST_ calculated between island and mainland populations was 0.081. It was on average 0.003 between close populations within the mainland and 0.005 within the island. *F*_ST_ was low, on average 0.003, for each of the four pairs of D- and E-populations, ranging from 0.0006 between B6D and B6E to 0.0079 between B4D and B4E. The *F*_ST_ estimated on the Z chromosome were on average 1.52 times higher than on the autosomes; 1.62 times higher when comparing populations from the mainland versus the Corsica Island and 1.44 times higher when comparing neighboring populations in the island or in the mainland.

### Demographic history

When deciphering the historical demography of deciduous and evergreen population pairs, the hierarchical model choice procedure strongly rejected models of strict isolation (SI), ancient migration and of panmixia (PAN), which were associated with a posterior probability of 0 (Supplementary table 3). Instead, models with gene flow were highly supported. In three of the four pairs, the equilibrium model (EQ) received the highest posterior probability with P(EQ) = 0.99 and 0.95, 0,94 in the B6D vs B6E, B5D vs B5E and B4D vs B4E comparisons, respectively. In the A2D vs A2E comparisons, the best supported model was the isolation with migration model, P(IM) = 0,74 with the second best model being SC, with P(SC) = 0.23. Comparing the two models against each other while excluding all remaining models yield unambiguous support for IM, with P(IM) =0.99 (Supplementary Table S3). Across all models, comparisons with heterogeneous gene flow and heterogeneous effective population size were not supported, indicating that, if genetic barriers or linked selection were at play they could not be detected. Demographic parameters were estimated for each pair of populations under the best model (Supplementary table 4). Posterior distributions were well differentiated from their prior indicating that estimated parameters were confidently estimated (Supplementary Figure 2). Effective population size (*N*_*e*_) where slightly higher in deciduous than in evergreen habitats (*Ne*1 in Table 2, Figure 3A, Supplementary table 4) and tended to be higher on the mainland than on the island. In a number of comparisons, our migration estimates reached the prior upper bounds. Therefore, we ran a new set of simulations with wider priors (Supplementary Note 1). Migration rates were not different between deciduous and evergreen habitats: in two instances, the rate of migration was higher from the deciduous to the evergreen and in two other instances, the reverse was true (Supplementary Figure 3).

**Figure 3.**
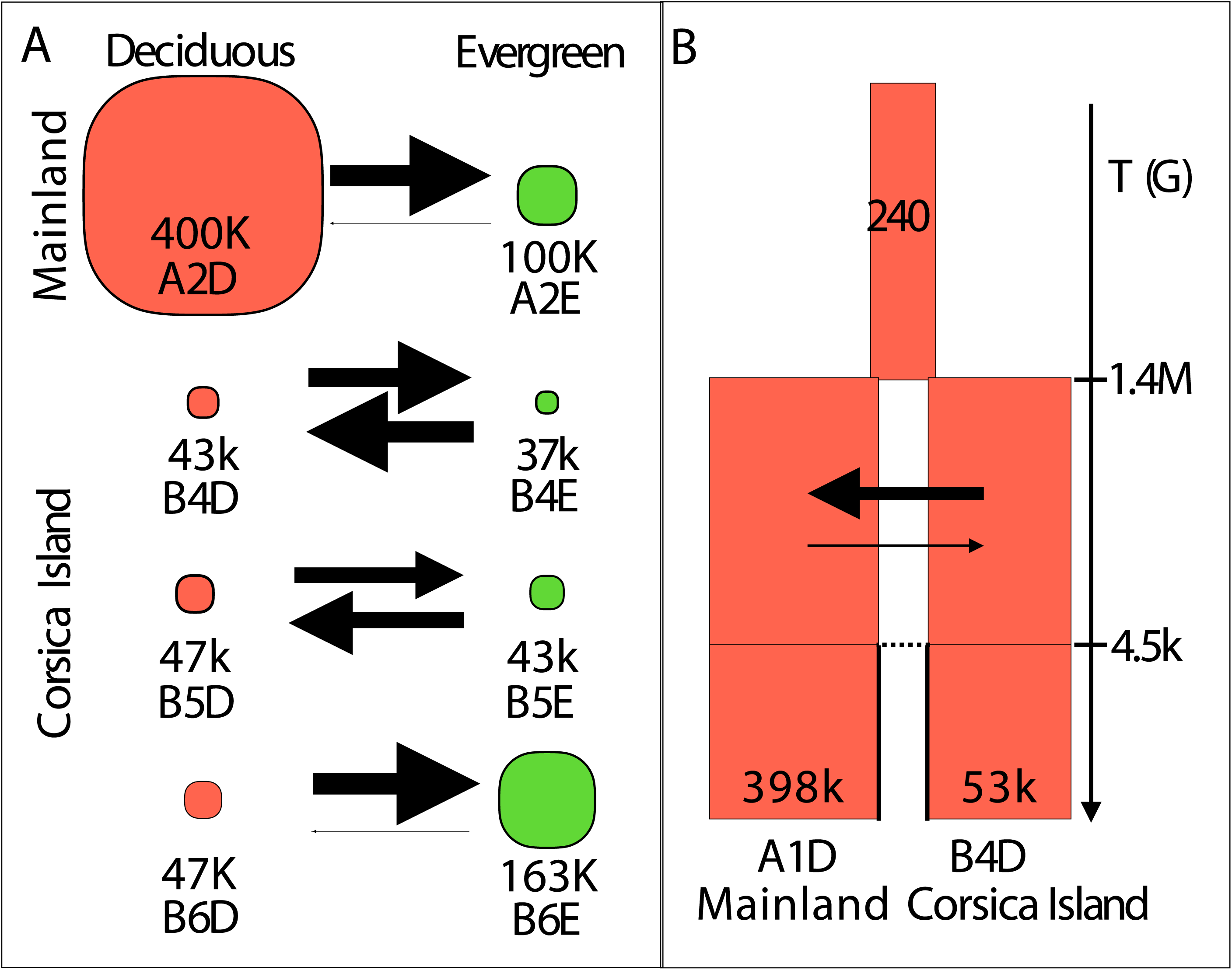
Demographic parameters. **A)** for each of the four pairs of blue tit populations in deciduous and evergreen habitats, estimated using equilibrium models, and **B)** for the divergence between populations on the mainland and Corsica, estimated using an ancient migration model. In panel A, circle size is proportional to effective population size (Ne) and arrow width is proportional to migration rate. In panel B, rectangle width is proportional to log10 (effective population size), arrow width is proportional to migration rate, and split time and time of ancient migration are indicated in number of generations.

Deciphering the historical demography between Corsica and the mainland, the ancient migration models (AM) strongly outperformed the strict isolation models (SI), with p(AM) = 0.99 (Supplementary table 5). For the AM model, the simplest model without linked selection or barriers to gene flow was the best supported, with p = 0.76. Parameters estimates revealed posterior generally well differentiated from the prior providing increased confidence (Supplementary Figure 4). Effective population size of the mainland was 397,685 [CI = 244,375 -- 600,105], around 7.5 times greater than the one of the island, 52,990 [CI = 33,065 -- 78,475] (Figure 3B, Ne1 in Table 2, Supplementary table 6). Our analysis also indicated a strong population size change during the process of population isolation, since the ancestral Ne was estimated at only 240 [CI = 65 -- 685]. Split time was estimated at 1,417,480 [CI = 740,000 -- 2,394,000] generations ago, hence around 3.2M years ago (assuming a 2.3 years generation time (Charmantier et al. 2004)) and gene flow subsequently stopped 4540 [CI = 2500 -- 6360] generations ago, hence around 10,000 years ago, corresponding closely with the end of the last glacial maximum. The ancestral migration rate between Corsica and the mainland, from the split in two populations to the end of the latest gene flow between these populations, was inferred asymmetric, with a 5 times larger migration rate from the island to the mainland (*m ∼ 1*.*9e-4*) than the other way around (*m ∼ 3*.*8e-5*). However, this asymmetry did not generate a strong difference in gene flow between the two groups, given the lower effective population size on the island compared to the mainland, (*i*.*e*. respective number of migrants of 10 and 15 from Corsica to the mainland and in the reverse direction, with overlapping credible intervals, see details in Supplementary table 6).

**Figure 4.**
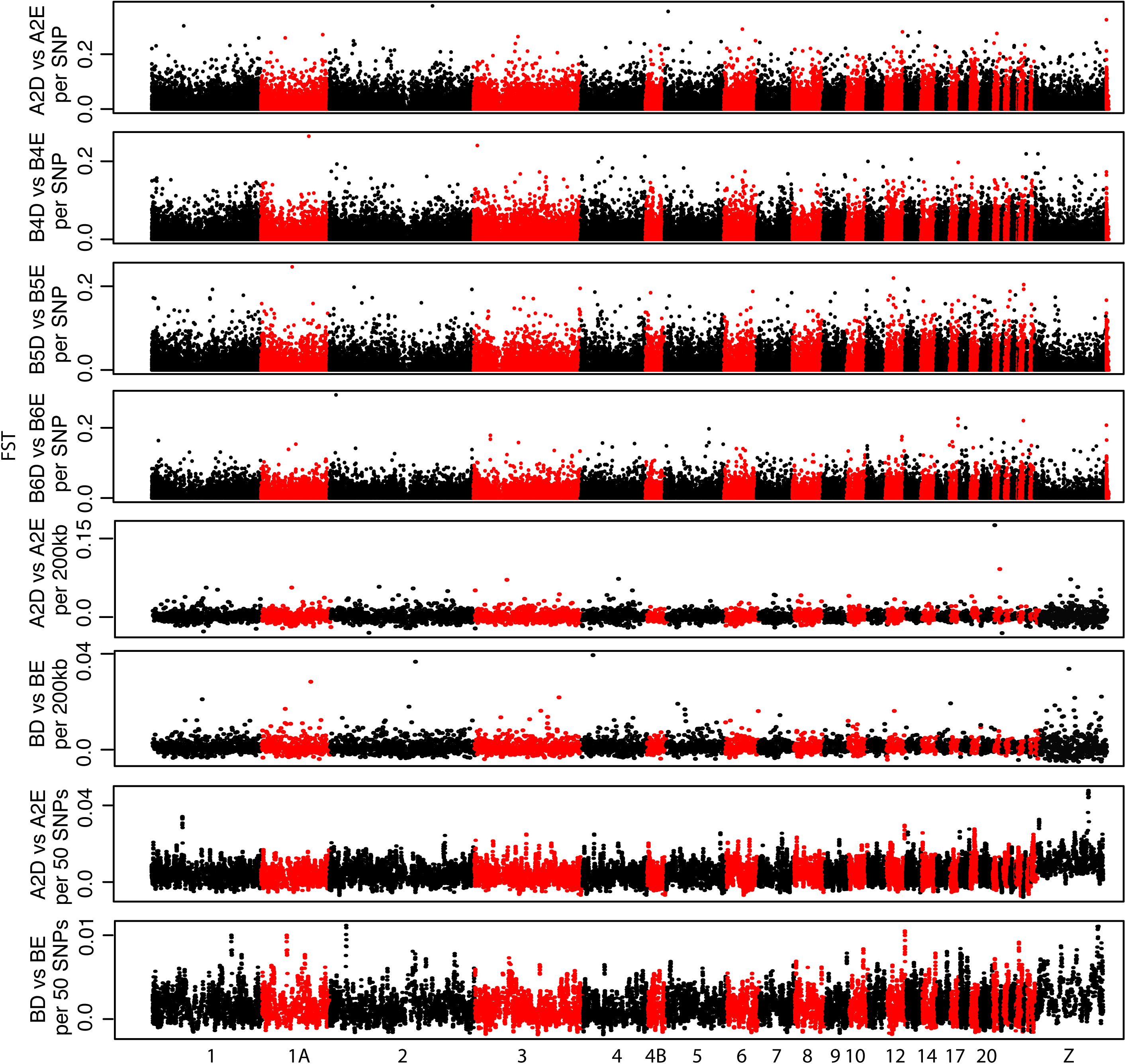
*F*_ST_ Manhattan plots between the 4 population pairs in deciduous and evergreen forests. *F*_ST_ are given either by SNP (four first graphs), by 200kb windows (graphs 5 & 6) or by 50 SNPs windows (graphs 7 & 8). For graphical simplicity, the 3 pairs of populations in Corsica were grouped for the sliding window *F*_ST_ Manhattan plots (BD *vs* BE). Dots alternate colors between chromosomes. *F*_ST_ for unplaced loci are shown at the end of each of the first four Manhattan plots.

### Genetic diversity

The mainland populations displayed significantly higher patterns of observed heterozygosity (on average 0.169, table 2) than Island populations (on average 0.147), all t-tests’ p-values < 0.001. On the contrary there was no significant difference of observed heterozygosity at smaller spatial scale among populations within the island (either same or different tree phenology, E- and D-) or within the mainland (all t-tests’ p-value > 0.05). More SNPs were polymorphic in mainland populations (77,306 to 80,616 SNPs for a MAF > 0.05) than in island populations (62,728 to 65,876 SNPs for a MAF > 0.05). The MAF spectrum showed enrichment of variants with smaller frequencies in mainland populations versus island populations (Supplementary Figure 5A). LD decayed rapidly in the first 5kb and was lower in populations from the mainland (especially A1D) compared to island populations (Supplementary Figure 5B). This pattern of rapid LD decay was similar between chromosomes (see for example chromosomes 1, 2 and Z, Supplementary Figure 6). Contemporary *N*_*e*_ inferred from LD varied from 142 (in A2E) to 355 (in A1D) (Ne2 in Table 2, Supplementary Figure 5C). *N*_*e*_ values were rather similar within pairs of E- and D-populations, although Ne was in average smaller in evergreen populations (245 in average) than in deciduous ones (261 in average). The four largest *N*_*e*_ (with an average of 318) were found for the four populations monitored on the long term in forest in which hundreds of artificial nest boxes have been installed for several decades (the *N*_*e*_ in the 6 other populations was in average of 210). In the entire dataset, 2599 SNPs were identified as non-synonymous while 6751 SNPs were identified as synonymous. We observed 1.18 times lower allele frequencies for non-synonymous variants (average MAF = 0.09) than for synonymous ones (average MAF = 0.11) (t.test p-value < 0.000001). This overall lower frequency of non-synonymous compared to synonymous mutations was similar between populations from Corsica (1.19) and the mainland (1.16), as well as between deciduous (1.18) and evergreen (1.19) habitats (See supplementary table 7). The Z chromosome harbored a proportion of non-synonymous mutations 1.83 times higher than the average for the autosomes. Finally, we observed significantly slightly longer ROH on Corsica compared to mainland (mean _island_ = 44,769 kb *vs* mean _mainland_ = 41,586 kb, Wilcoxon rank sum test W = 32,695, p <2e-16, Supplementary Figure 7&8). We also observed significant differences in the count of ROH among populations from the mainland versus those from the island but not among populations within the mainland or the island (ANOVA p <2e-16; Supplementary Table 8 for Tukey HSD test).

**Figure 5.**
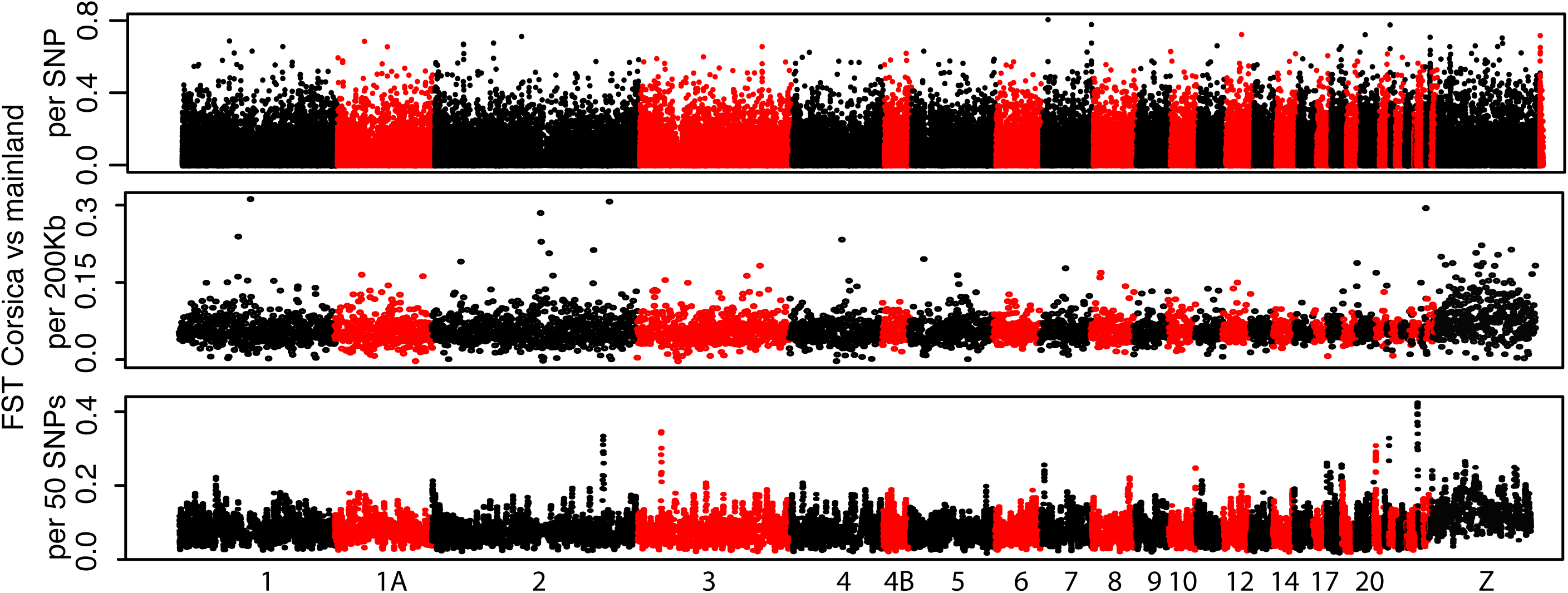
*F*_ST_ Manhattan plots between Corsica and the mainland. *F*_ST_ are given either by SNP, by 200kb windows or by 50 SNPs windows. Dots alternate colors between chromosomes. *F*_ST_ for unplaced loci are shown at the end of the first Manhattan plot.

**Figure 6.**
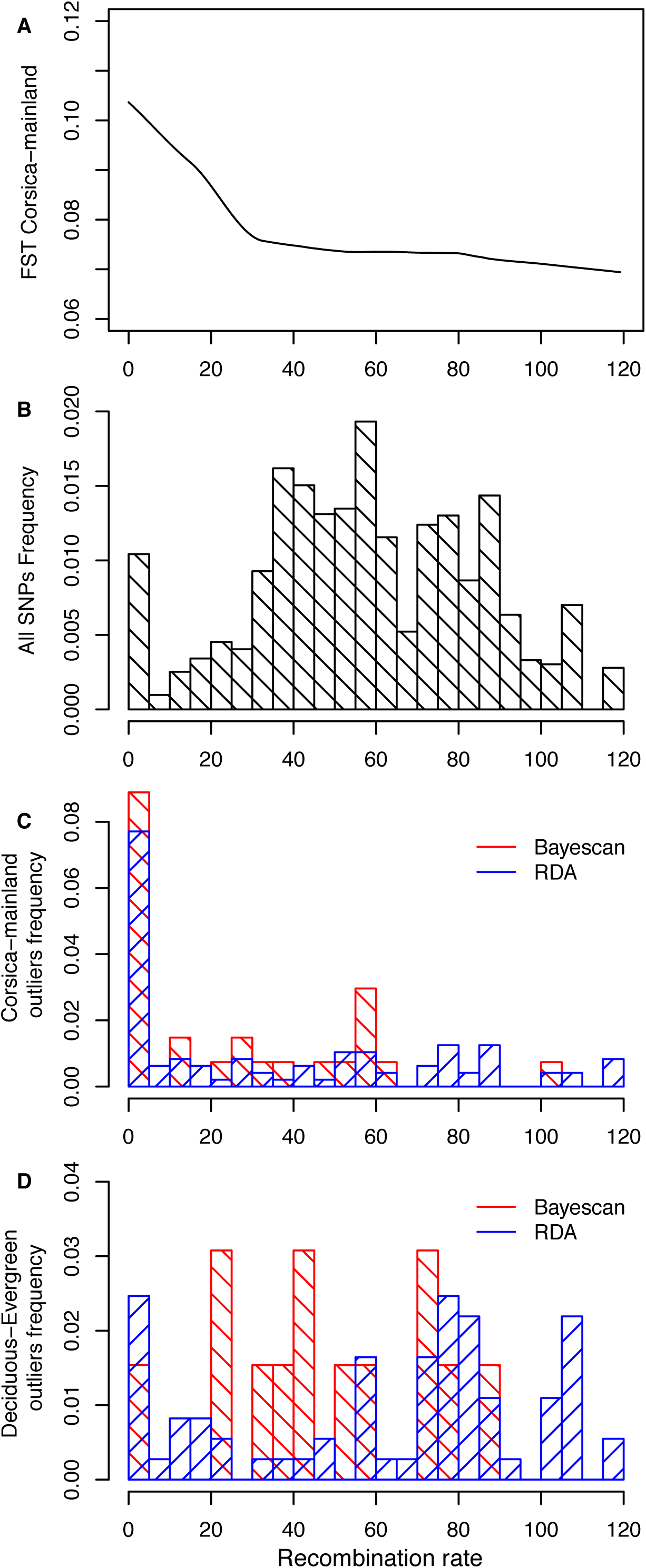
Relationship between recombination rate and divergence. **A)** Correlation between local recombination rate and *F*_ST_ between populations on the mainland and in Corsica; **B)** SNP frequency per class of recombination rate; **C)** Outlier SNP frequency for the mainland-Corsica comparison, per class of recombination rate; **D)** Outlier SNP frequency for the deciduous-evergreen comparisons, per class of recombination rate. In panels C & D, outliers are given for the Bayescan and the RDA tests.

**Figure 7.**
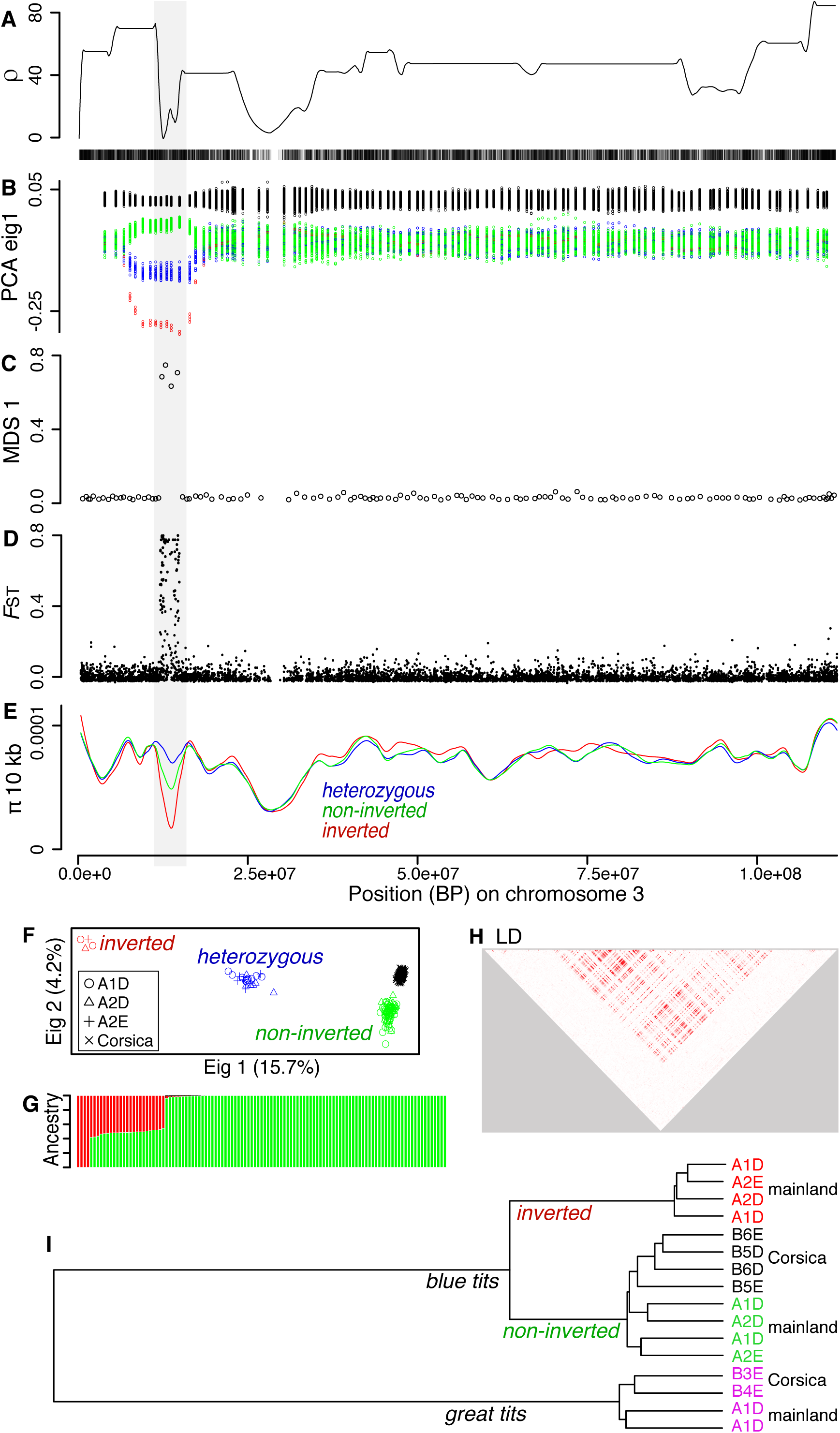
Detection and characteristics of a putative inversion on chromosome 3. A) Recombination rate and SNP density along chromosome 3; B) individual values for eigenvector 1 of a PCA of all the blue tit individuals from Corsica and the mainland; C) MDS 1 values from Lostruct analysis; D) *F*_ST_ between inverted, non-inverted and heterozygous individuals from the mainland; E) Nucleotide diversity for inverted (in red), non-inverted (in green) and heterozygous individuals (in blue) from the mainland; F) PCA results for the region of the putative inversion for all the blue tit individuals; G) Mainland blue tits’ admixture proportions for K = 2 for the region of the putative inversion (individual ordered by decreasing ancestry value for the inverted cluster); H) Linkage disequilibrium for SNPs found in the putative inversion (plus 1 Mb each side of the inversion), for mainland blue tit; I) genetic distances at the inversion, between 4 inverted blue tits, 8 non-inverted blue tits, and 4 great tits (sites of capture are indicated for each individual).

### Footprints of selection

Manhattan plots of *F*_ST_ per SNP revealed no clear evidence for high peaks of *F*_ST_ for the five comparisons considered (First panels of Figures 4 & 5). Throughout the five Bayescan tests (Corsica vs mainland populations and the 4 D-4 population pairs), we identified 40 SNPs with a log10(BF) > 0 among which 18 SNPs had a log10(BF) > 1 (Supplementary table 9, Supplementary Figure 9). None of these SNPs were detected twice among the different tests conducted with Bayescan. Among these 40 SNPs, 27 were found for the Corsica vs mainland test, 2 were found for A2D vs A2E, 9 were found for B4D vs B4E, 0 were found for B5 vs B5E and 2 were found for B6D vs B6E. Among these 40 SNPs, 19 genes were identified (Supplementary table 9). Sliding windows of F_ST_ along the genome showed a few outlier windows of modest intensities (last panels of Figure 4 & 5), with almost no parallelism between the tests (Supplementary Table 10). Expectedly, outlier windows found for the Corsica *vs* mainland comparison depicted larger *F*_ST_ than for deciduous vs evergreen comparisons. The outlier *F*_ST_ windows falling in the top 1% of the *F*_ST_ windows distribution are reported in Supplementary Table 10. Using the RDA to identify outlier SNPs with extreme loading values to axes, we identified 229 outlier SNPs associated to habitat type (deciduous vs evergreen), and 227 SNPs associated to geography (longitude & latitude, essentially representing a Corsica-Mainland comparison) (Supplementary table 11, Supplementary Figure 10). For the deciduous vs evergreen test, only one of the RDA outliers was also found outlier in Bayescan tests. For the Corsica-mainland test, 8 of the RDA outliers were also found outliers in the Bayescan test. Genes in which the outliers were found are reported in the supplementary table 13.

### Variation of genomic differentiation with recombination rate

*F*_ST_ between mainland and Corsican populations was negatively correlated to recombination rate (Figure 6A, linear model p < 2e-16; Supplementary Figure 11). In contrast, *F*_ST_ between each pair of deciduous and evergreen populations was not correlated to recombination rate. *F*_ST_ outlier SNPs between Corsica and the mainland populations and identified by Bayescan or by the RDA were more often found in regions of low recombination (χ2 test p-values < 0.01, Figure 6C) than observed for the entire SNPs (Figure 6B). Average recombination rate was on average twice lower for these F_ST_ outlier SNPs compared to the rest of the SNPs (t-test p-values < 1e-6). In contrast, *F*_ST_ outlier SNPs between deciduous and evergreen populations were not preferentially found in regions of low recombination (Figure 6D).

### Genomic inversions

We detected one putative inversion on chromosome 3, spanning 2.8Mb, from position 11,838,789 to 14,661,550, and containing 390 SNPs. The estimated recombination rate was on average 9 times lower (t-test: p<2.2e-16) in this region of the chromosome compared to the rest of the chromosome (Figure 7A). However, the SNP density did not appear reduced at the location of this putative inversion (Figure 7A), suggesting a recent drop of recombination (*i*.*e*. dissimilar to what is expected in a peri-centromeric location). We did not find evidence for an increase of read depth at specific loci in the putative inversion or its vicinity in inverted individuals compared to normal ones. We did however notice the absence (read depth = 0) of 3 loci from the inversion, in the four inverted individuals. The sliding PCA detected either 2 groups (individuals from Corsica and from mainland) outside of the putative inversion, or 4 groups at the putative inversion location, separating individuals from Corsica and 3 groups for mainland individuals (Figure 7B). We assumed that the inverted segment was the one with the lower frequency on the mainland and that was absent from Corsica. The Lostruct analysis confirmed for this same region the presence of four successive blocks of 100 SNPs highly discriminating individuals (Figure 7C). The average *F*_ST_ between the three groups of mainland individuals was much higher (t-test: p<2.2e-16, Figure 7D) for SNPs in the putative inversion (*F*_ST_ = 0.26) than for SNPs outside of this region (*F*_ST_ = 5.8e-4). π was lower (t-test: p=1.9e-5, Figure 7E) for SNPs located in the putative inversion for inverted homozygous individuals (π = 2.5e-5) compared to non-inverted homozygous individuals (π = 5.3e-5). Both inverted and non-inverted homozygous individuals had lower π at this inversion compared to heterozygous individuals at this region (π = 7.6e-5) and compared to the three types of individuals for the rest of the chromosome 3 (π ranging from 7.4e-5 to 7.6e-5). The PCA (Figure 7F) and the admixture analysis (Figure 7G) using the SNPs from the inversion clearly delineated inverted, non-inverted and heterozygous individuals, with heterozygous individuals falling at intermediate eigenvalue between the two categories of homozygous individuals. LD was higher (Figure 7H) for the region of the putative inversion (Little LD was found when only homozygous non-inverted individuals were kept in the analysis). For the region of the inversion, Dxy was 0.0020 between normal blue tits from the mainland and from Corsica as well as between great tits from the mainland and from Corsica. Dxy was 0.0044 between inverted and normal blue tits from either the mainland or from Corsica, and ranged from 0.0149 to 0.0152 for interspecific comparisons. Using the formula T = Dxy / 2µ, the inversion likely appeared approximately 220,000 generations ago (around 506,000 years ago). The tree of genetic distance illustrated the divergence of inverted blue tits from non-inverted blue tits from both mainland and Corsica (Figure 7i). This region contains 21 genes listed in supplementary table 12. Out of 113 individuals from the mainland, we found 4 inverted homozygous, 23 heterozygous, and 86 non-inverted homozygous, hence we observed no deviation from the Hardy-Weinberg equilibrium (χ2 test p-value > 0.1). The percentage of each genotype was similar across all mainland locations and both habitat types (the inverted segment was at 11% in A1D, 16% in A2D, 17% in A2E, and 14% across these 3 locations).

### Gene ontology

None of the gene lists gathered with the different tests (each outlier test among deciduous and evergreen environment and between Corsica and the mainland and the inversion test) yielded any significantly enriched GO term after correction multiple testing (Supplementary table 14). The most promising GO (uncorrected p.value <0.0025) included: i) for the deciduous-evergreen outlier tests, GO:0060385, axonogenesis involved in innervation; GO:2001013, epithelial cell proliferation; GO:0007194, negative regulation of adenylate cyclase; GO:0090647, modulation of age-related behavioral; GO:1901379, regulation of potassium ion transmembrane transport, ii) for the Corsica-Mainland outlier tests, GO:0008344, adult locomotory behavior; GO:2001224, positive regulation of neuron migration, iii) for the inversion, GO:0016446, somatic hypermutation of immunoglobulin; GO:0045910, negative regulation of DNA recombination; GO:0006298, mismatch repair; GO:0008340, determination of adult lifespan.

## Discussion

In this study, we investigated demographic history and genome wide patterns of genetic diversity and differentiation between several populations of blue tit presenting strong phenotypic differences between deciduous *versus* evergreen forest habitats as well as insular versus mainland contexts. Between populations in deciduous and evergreen forest habitats, demographic modelling showed large gene flow and large effective population sizes (Ne), explaining the low genetic differentiation between these populations. Demographic modeling also revealed that each pair of deciduous and evergreen populations most likely never diverged and maintained high connectivity through migration. We identified slight and mostly unrepeated footprints of divergent selection across these evergreen and deciduous population pairs, which is consistent with their demography and the likely polygenic nature of several traits implicated in their local adaptation. In both insular and mainland populations, we found large Ne, although smaller for insular populations than mainland ones, resulting in different distributions and lengths of ROH. Gene flow between Corsica and the mainland most likely stopped at the end of the last glaciation. Both large Ne and gene flow until the end of the last glaciation hence explained moderate genome-wide differentiation. We identified several genomic footprints of selection, enriched in regions of low recombination only in the case of the mainland/island divergence. Finally, we identified a putative genomic inversion spanning 2.8Mb, polymorphic in mainland populations only. We discuss these results in the context of the vast scientific knowledge acquired on these blue tit populations during the last four decades and more generally in the context of genomics of adaptation with gene flow.

### Divergence between populations in deciduous *versus* evergreen habitats

Although we found a significant effect of habitat on genetic structure, the genetic differentiation between neighbouring deciduous and evergreen populations was low (*F*_ST_ ranging from 0.0006 to 0.0079). This result is in line with the primary observations realized earlier on a smaller set of populations (Porlier et al. 2012b; Szulkin et al. 2016). Accordingly, we found high gene flow from deciduous to evergreen populations. Yet, this quantification of high gene flow and low genetic structure contrasted with the demographic knowledge collected on blue tit. Indeed, demographic studies suggested restricted dispersal between these populations, with 4 dispersal events observed between B4D and B4E (5.6 km apart) and none between the B3E and either B4D or B4E (24.1 km), among a total of 2788 males, 2672 females and 25,158 nestling ringed in the three main Corsican sites between 1976 and 2018, with a mean recruitment rate of 6% (Charmantier, com pers). Our interpretation of this contrast between gene flow estimations gathered from population genomic *versus* recapture data is quadruple. First, dispersal estimation on the field using capture-mark-recapture is very challenging and may require more data than currently collected, to detect rare dispersal events, even though these affect population genetic parameters. Moreover, since natal dispersal in blue tit classically ranges between 330 m and 4 km (see (Tufto et al. 2005; Ortego et al. 2011)), the long-term monitoring sites in Corsica equipped with nest-boxes (black circled dots in Figure 1C) are not ideally spaced to identify the origin of immigrants and the destination of emigrants, and only a small fraction of the landscape favourable to blue tit breeding is covered. Second, only a few migrants are sufficient to decrease the genetic distance between populations, measured through the *F*_ST_ (Marko & Hart 2011; Cayuela et al. 2018). In that regard, our results may be compatible with the few dispersal events recorded throughout the years. Third, it is important to note that the number of migrants estimated using demographic analyses represents an average over historical time scales on the order of Ne generations and may have varied widely during contemporary times. Fourth, the large effective population size found both using coalescence and a LD method might be explained by the existence of large “metapopulations” connected by high gene flow and such large Ne might largely contribute to limit the genetic divergence between populations.

Ne was on average slightly larger in deciduous than in evergreen populations. This could be explained by the higher productivity of deciduous forests compared to evergreen forests, resulting in larger clutches and more fledglings in deciduous habitats (Table 1 in Charmantier et al 2016). However, the very high gene flow between deciduous and evergreen populations and the low genetic differentiation between these populations limits further interpretations. In addition, populations monitored on the long term, for which hundreds of artificial nest boxes have been installed, tended to have larger Ne than non-monitored populations. Breeding density in the nest-box areas studied (Figure 1A) was around 1 to 1.3 pairs per ha (Blondel et al 2006), which is most probably 3 to 5 times higher than natural densities for blue tit when these secondary hole-nesting birds rely on natural cavities only. It is hence possible that the recent availability of nest boxes locally increased productivity, effective population size and heterozygosity.

Demographic modeling indicated that secondary gene flow between previously isolated populations, which could have eroded most genome-wide differentiation outside regions implicated in local adaptation and/or acting as reproductive barriers in particular in low recombining areas (Bierne et al. 2013), was very unlikely. Secondary contacts were strongly rejected in favor of an island model supporting ongoing gene-flow without divergence (EQ model) in 3 out of 4 population pairs, the last one supporting an isolation with migration model (IM). As is often the case, our demographic estimates should be interpreted cautiously, particularly in the absence of a well-documented mutation rate in blue tit. The relatively narrow credible intervals around parameter estimates nevertheless provide confidence in their biological relevance. These results suggest that the focal population pairs occupying deciduous/evergreen habitats have been continuously connected by relatively high gene flow. Hence, genetic divergence between populations in each pair did not originate or build up on pre-existing genetic divergence accumulated during an allopatric phase but rather is supported as a case of genuine recent ecological divergence (Pinho & Hey 2010; Wang & Bradburd 2014).

We did not find strong footprints of divergent selection between any of the four pairs of deciduous and evergreen populations (Figure 4). Moreover, the observed outliers were not found repeatedly across the 4 deciduous-evergreen comparisons but rather each outlier was found only once, and they were not enriched in regions of low recombination. First, this result is in line with the high migration rates observed. High gene flow indeed most likely limits the potential for local adaptation since it limits the accumulation of allelic differentiation, even with high selection coefficients (Lenormand 2002). Second, this pattern is consistent with the best demographic model being the equilibrium model and not a secondary contact. Indeed, the later would more often create repeated and strong outliers located in regions implicated in reproductive isolation between divergent populations (eg (Rougemont et al. 2017)). Third, this pattern is congruent with a model of local polygenic adaptation involving a transient genetic architecture with multiple alleles of small effects underlying (multiple) quantitative characters (Yeaman & Whitlock 2011; Yeaman 2015) and displaying low F_ST_ among loci under divergent selection. Fourth, most documented traits that are involved in the adaptation of blue tit to the deciduous *versus* evergreen habitat types, such as clutch size or laying date (Lambrechts et al. 1997; Blondel et al. 1998) are quantitative and most likely rely on a polygenic architecture with alleles of small effects (see e.g. (Santure et al. 2013)). However, some statistical issues may limit the interpretation of this result. First, currently available genome-scan methods to detect alleles of small effect are statistically limited, especially when applied to relatively small datasets (Rockman 2012; Hoban *et al*. 2016). Second, a potential lack of power in detecting outliers could arise from the combination of the use of RADsequencing and a rapid LD decay along the genome, that may result in too low resolution especially in regions with larger recombination rates (Lowry et al. 2017). Further analyses of the potential outliers found here, as well as a new analysis with higher marker density and more individuals, are needed to better document and discuss the potential genes and biological processes implicated. Particularly, quantitative genomics will be useful to establish links between the level of divergent selection on these putative targets and their effect size on phenotypic trait variation (Stinchcombe & Hoekstra 2007; Gagnaire & Gaggiotti 2016). This would notably reveal whether genes under divergent selection are also responsible for the observed phenotypic variation, and contribute to assessing of how much of phenotypic variation is adaptive.

Last, we searched for inversions potentially associated with phenotypic variation and/or segregating in both habitats. We report multiple evidences (Figure 6) for a putative inversion on chromosome 3 in mainland populations. However, the proportion of inversions in the genomes of deciduous-*versus* evergreen-breeding birds did not differ, which most probably precludes a putative role of the inversion in local adaptation in these habitats. Besides, this polymorphism followed the Hardy-Weinberg equilibrium and we found 4 homozygous inverted individuals, suggesting this inversion does not involve an accumulation of lethal recessive deleterious alleles (Jay et al. 2019). The functional consequences of this inversion on variation in life history traits in blue tit call for further investigations, as achieved in other songbirds (Tuttle et al. 2016; Knief et al. 2017; Kim et al. 2017). In order to genotype this inversion in more birds and to link it to putative phenotypic variation, we could use a PCR-RFLP approach (da Silva et al. 2019).

### Divergence between blue tit populations in mainland *versus* insular contexts

Our demographic modelling approach of Corsican and mainland populations revealed that a model with ancient migration was the most probable, which is coherent with the history previously reconstructed for the blue tit complex. Split time of the ancestral populations in two mainland *versus* island populations connected by gene flow (∼3M years) was relatively coherent with the diversification time found in the literature for the entire blue tit complex (5M (Illera et al. 2011)). Gene flow between the island and the mainland likely stopped around 10,000 years, which is compatible with a gene flow break due to the rise of the flooded stretch between Corsica and the mainland during ice melting and the sea level rise after the last glaciation, during from 17,000 to 5,000 years ago (Lambeck & Bard 2000; Jouet et al. 2006). Both the signal of population expansion during the diversification period, and the gene flow five times larger from the island to the mainland than from the mainland to the island, were coherent with a suspected recolonisation of the mainland from the islands, as proposed by Illera et al. (2011) based on analyses of nuclear and mitochondrial DNA sequences. The larger effective population size on the mainland than on the island may be due to multiple sources of colonisation on the mainland (Taberlet et al. 1992) and to the much larger mainland area and hence metapopulation size compared to the island. Expanding this study with a sampling from the South-East of Europe, one from Sardinia and one from the Iberian Peninsula, could help determine whether the large size estimated for the mainland meta-population studied here might be explained by a mixed recolonisation from these refugium (Kvist et al. 1999; 2004) and whether insular populations consistently have reduced effective population size. It would also be interesting to explore a model allowing for multiple cycles of isolations and contacts that could have resulted from successive glacial cycles (Hewitt 2004).

Given the demographic parameters inferred, it is not surprising that the genetic differentiation between populations from the mainland and the island was moderate (*F*_ST_ = 0.08). However, this differentiation value contrasted with the much larger divergence at mitochondrial DNA observed for these populations (OST = 0.67 between A1D and B4D (Kvist et al. 2004)). This discrepancy between whole nuclear genome and mitochondrial estimates of differentiation could first be explained by the typically four times smaller Ne in mitochondrial DNA compared to nuclear DNA in diploid organisms, resulting in larger drift and divergence (Smith & Klicka 2013). Second, accumulation of cytonuclear incompatibilities could limit its introgression (Burton & Barreto 2012). Alternatively, gene flow may have been greater for males than females during the colonisation process from Corsica to the mainland, at the end of the last glacial period. However, this hypothesis is not supported by the general observation that females disperse on average further away than males and that rare long distance dispersals are also achieved by females (Tufto et al. 2005). Inspecting mitochondrial DNA variation for the birds studied here would be very useful to compare mitochondrial differentiation in our sample to the previous analysis which also included Corsican birds (Kvist et al. 2004). In any case, one should bear in mind that our results stem from integrating coalescent patterns observed across thousands of loci, therefore providing increased resolution to investigate the determinants of demographic divergence compared to approaches based on fewer mitochondrial and nuclear loci.

The contemporary demography inferred between the populations from Corsica and mainland France, especially the absence of gene flow for approximately 4500 generations and the large effective population size, likely provides a suitable context for the build-up of local adaptation. Moreover, the level of divergence (*F*_ST_ = 0.08) was probably low enough to detect divergent outlier loci potentially implicated in local adaptation. We identified several outlier SNPs and outlier 200 kb windows showing elevated differentiation, more often in regions of lower recombination than elsewhere in the genome (contrary to the outliers found for the deciduous vs evergreen populations). Furthermore, *F*_ST_ was on average higher in these regions of low recombination compared to elsewhere in the genome (which was not the case for deciduous vs evergreen populations). These patterns of increased divergence in regions with low recombination between divergent populations are commonly observed in other species (Nachman & Payseur 2011; Cruickshank & Hahn 2014; Gagnaire et al. 2018), including other bird species (Burri et al. 2015; Spurgin et al. 2019). It is well documented that such increased differentiation in regions with low recombination is not necessarily due to positive selection, or at least not alone, and that it is largely influenced by the effect of recombination in interaction with background selection (Charlesworth et al. 1993; Burri *et al*. 2015; Perrier & Charmantier 2019). Moreover, a uniform *F*_ST_ sliding window, sized in Kb, is expected to dilute signatures of selection in regions of the genome where LD is fast decaying while it is expected to over-represent regions where LD decays much more slowly, hence increasing even more the potential false positives in regions with low recombination (Beissinger et al. 2015; Perrier & Charmantier 2019). Although perfectible, our sliding window approach using a SNP unit attempted to fix this issue and successfully captured several new outlier regions outside of deserts of recombination. This second window approach should however be improved, for example by estimating the local neutral enveloped, which computation should integrate local variations in LD, recombination, diversity, but also the demography of populations. As mentioned for the study of adaptive divergence between deciduous and evergreen populations, complementary analyses integrating both genome scans and quantitative genomics would improve our comprehension of the genomic and phenotypic divergence observed between Corsican and mainland blue tits (Stinchcombe & Hoekstra 2007; Gagnaire & Gaggiotti 2016).

While the putative genomic inversion on chromosome 3 was absent from Corsica and detected in mainland individuals, its level of divergence from the non-inverted sequence indicated that it was likely twice older than the beginning of divergence between blue tit populations from mainland and Corsica. This could first suggest that this polymorphism emerged in mainland blue tit populations and then did not introgress the Corsican populations, maybe due to a local disadvantage and/or genetic incompatibilities or simply due to drift coupled to little gene flow. A second hypothesis could be that this inversion was present in Corsican populations but had been purged out due to a local disadvantage. Such a disadvantage can be due to the typical accumulation of deleterious mutations in such non-recombining inverted sequences (*e*.*g*. Jay et al 2019). Lastly, it is possible that this inversion has been acquired in mainland populations only recently, after the last period of contact between mainland and Corsican populations, via gene flow from a distinct refugium with which connectivity could have been enhanced after deglaciation, hence after the stop of gene flow with Corsica. The age of this inversion, its origin, its biological effects and the potential accumulation of deleterious mutations need to be inferred more thoroughly *via* genotyping full inversion sequences from individuals in diverse locations and using advanced statistical methods (Lohse *et al*. 2015).

### Conclusion and perspectives

Our study demonstrated the usefulness of demographic modeling and of the analysis of the variation of genomic diversity and recombination along the genome to uncover the genetic determinants of local adaptation in a small passerine with a large distribution, and occupying different forest habitats. Especially, demographic modeling rejected the hypothesis of a secondary contact between deciduous and evergreen populations and favoured a situation with continuous gene flow. These results support the idea that blue tits have adapted to their habitats despite ongoing gene flow, while contextualising how large gene flow most probably constrained local adaptation (Lenormand 2002) and favoured its architecture based on alleles of small effects (Yeaman 2015). The genomic modelling also refined our knowledge about the divergence between insular and mainland meta-populations, that have been likely unconnected by gene flow for the last ten thousand years. We also verified the relationship between local recombination rate and differentiation, that should probably be integrated in genome scans looking for footprints of selection (Beissinger *et al*. 2015; Burri *et al*. 2015; Berner & Roesti 2017; Perrier & Charmantier 2019, Booker et al 2020). Future investigations will require increased sample sizes and marker density (Lotterhos & Whitlock 2015; Hoban *et al*. 2016) in order to better detect loci with small effects that contribute to the quantitative phenotypic variation and local adaptation in blue tit. Lastly, the putative inversion found here would need further analyses since this type of structural variation is often implicated in phenotypic variation (Kirkpatrick 2006; 2010; Stapley *et al*. 2017; Wellenreuther *et al*. 2019).

## Acknowledgments

We wish to thank the many people who helped on the blue tit long-term project. We particularly wish to thank Philippe Perret. We also wish to thank Samuel Perret, Christophe De Franceschi, Jacques Blondel, Marta Szulkin, Monica Arias, Patricia Sourouille, Annick Lucas & Boris Delahaie. This project was funded by the European Research Council (Starting grant ERC-2013-StG-337365-SHE to AC) and the OSU-OREME.

## Authors’ contributions

CP conceived the study, conducted field-work, lab analyses, bioinformatics and statistical analyses, interpreted the results and wrote the manuscript. QR conducted bioinformatics and statistical analyses (especially demographic analyses using ABC), interpreted the results and edited the manuscript. AC obtained the ERC grant, conceived the study, performed field-work, interpreted the results and edited the manuscript.

## Data accessibility

Individual sequences with metadata are available on NCBI with the SRA accession PRJNA630135. A filtered VCF file with metadata are available on Dryad with the doi:10.5061/dryad.x69p8czfg.

