## Supplementary figures and images for "Demographic history and genomics of local adaptation in blue tit populations"

### Supplementary Figure 1

Supplementary Figure 1. 5 PCA between i) all the individuals, ii) A2D vs A2E, iii) B4D vs B4E, iv) B5D vs B5E, v) B6D vs B6E

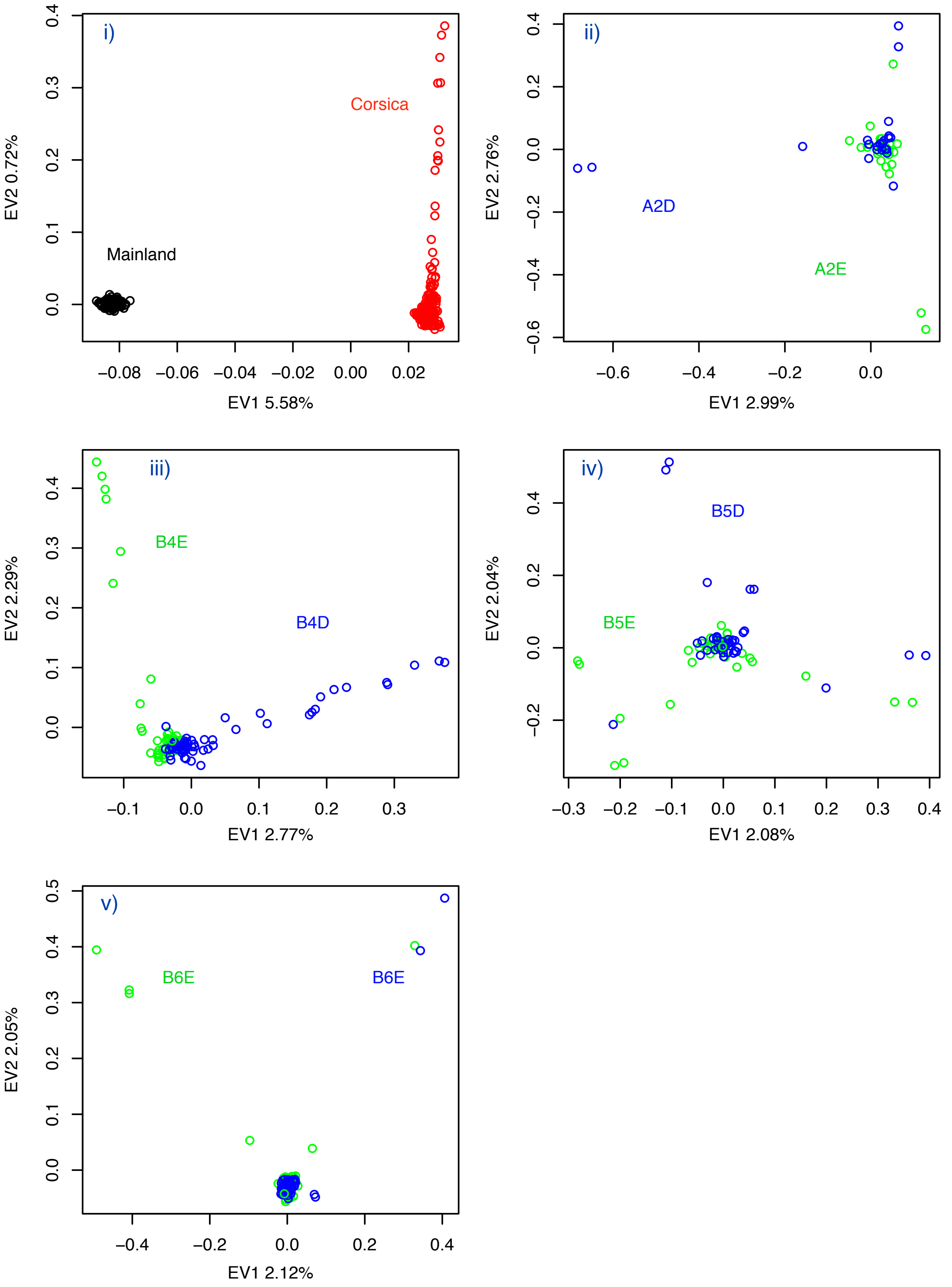

### Supplementary Figure 7

Supplementary Figure 7. Total length of ROH as a fonction of total number of ROH, per population.

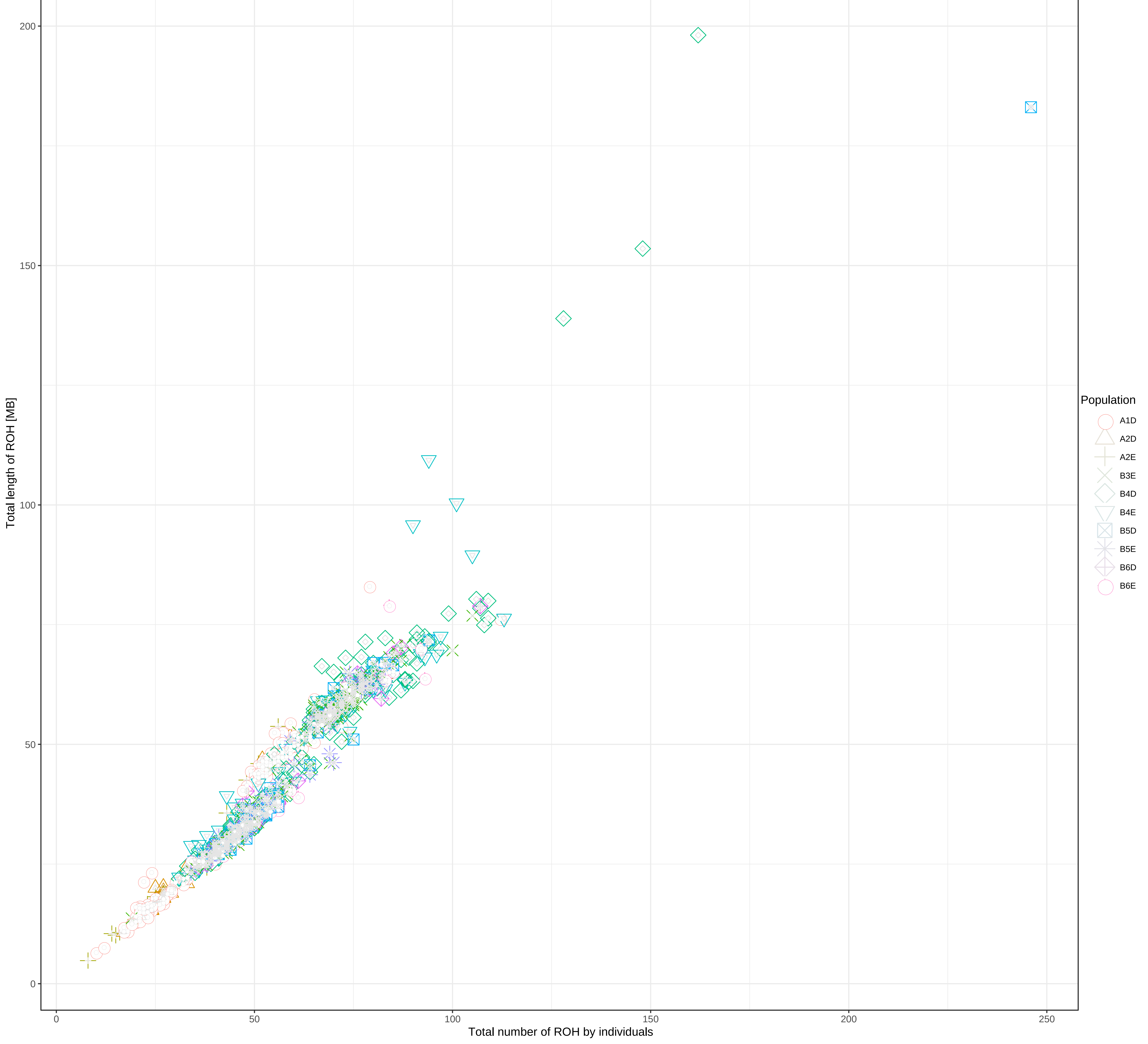

### Supplementary Figure 8

Supplementary Figure 8. Boxplot of ROH length (Kb) per population

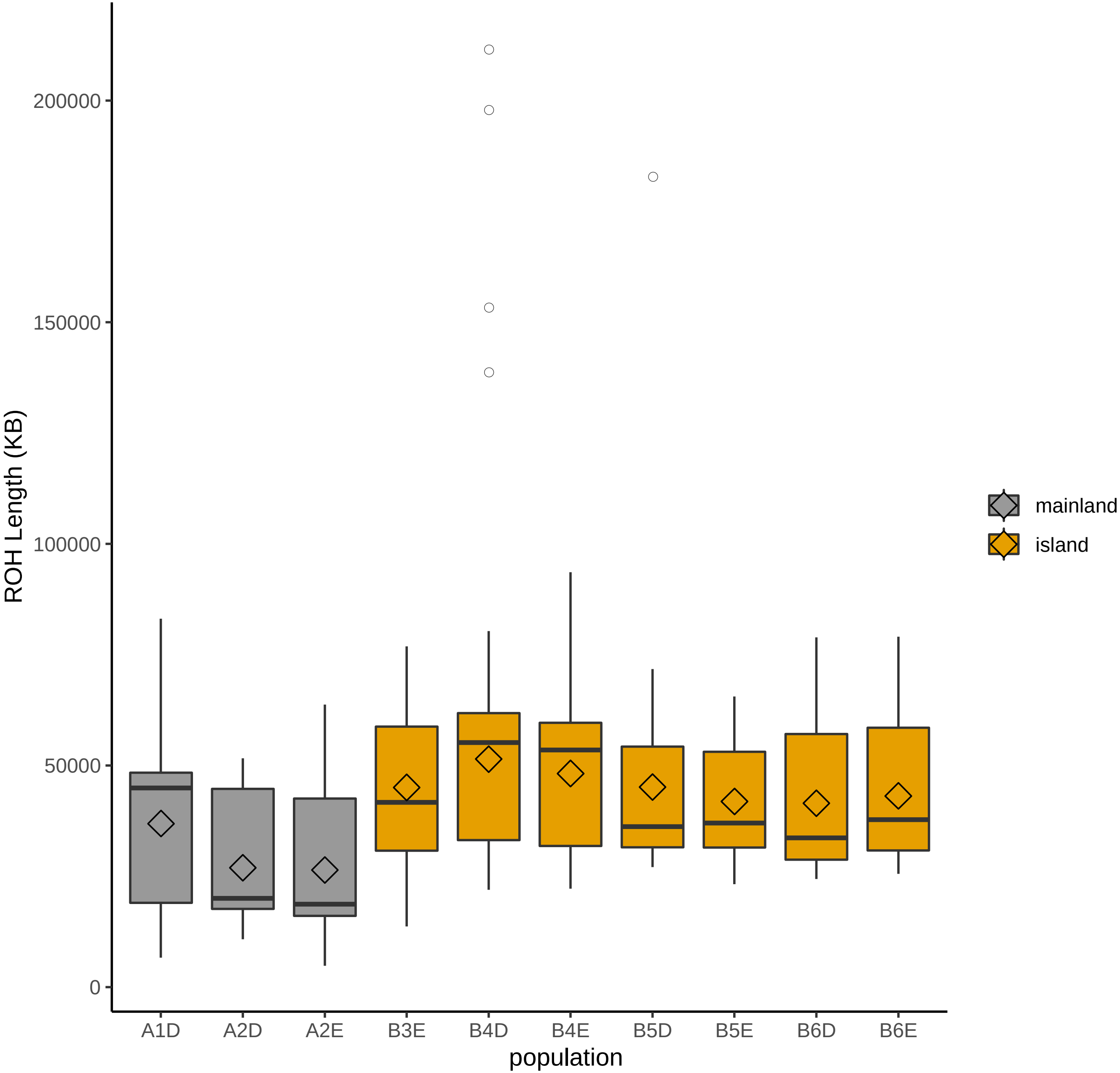

### Supplementary Figure 11

Supplementary Figure 11. Fst Corsica-Mainland as a function of recombination rate

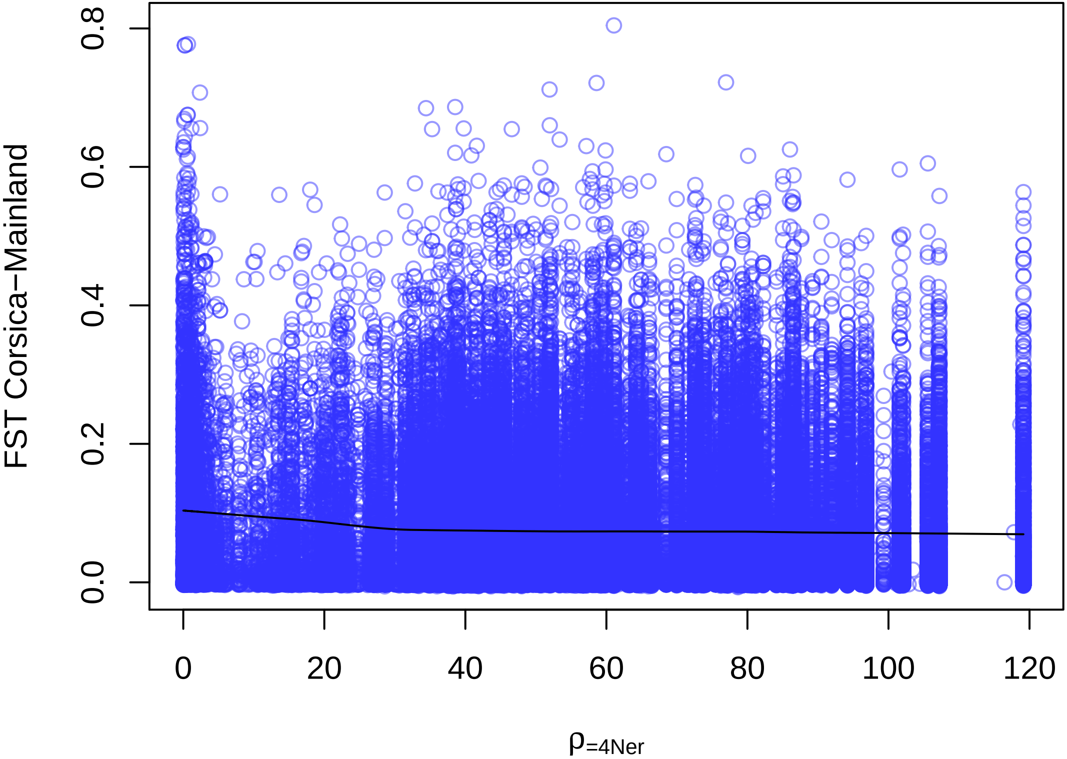
