## Supplementary Figure 2 for "Demographic history and genomics of local adaptation in blue tit populations"

**Supplementary Figure 2: Prior and posterior probabilities of estimated parameters for each pair of population.**

Theta1 = 4N1/µ (scale mutation rate for the population 1), Theta2 = 4N2/µ (scaled mutation rate for population 2). ThetaA = 4Nanc/µ/. (scaled mutation rate for the ancestral population. All theta are scale by the theta of the reference population (4*Nref*µ). Migration rate $M_{ij}$ = 4N * ref *$m_{ij}$. . Split time for the IM model is scaled by the 4Nref so that τ =Tsplit/4Nref.

#### A2DE – Best model = Isolation With Migration

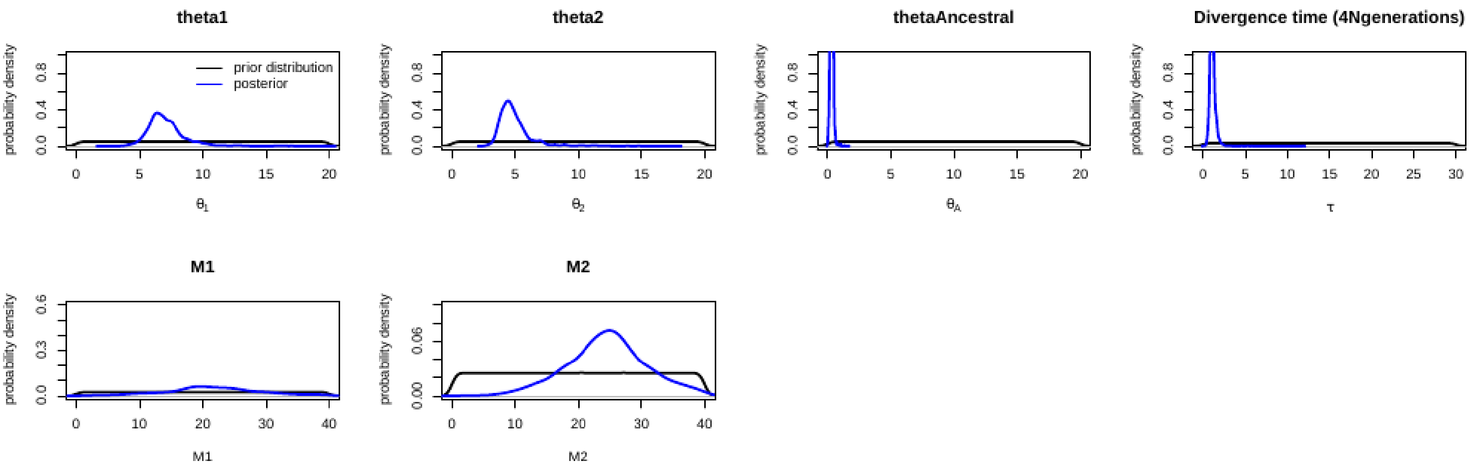


#### A2DE – Second best model:  = Equilibrium (Island model)

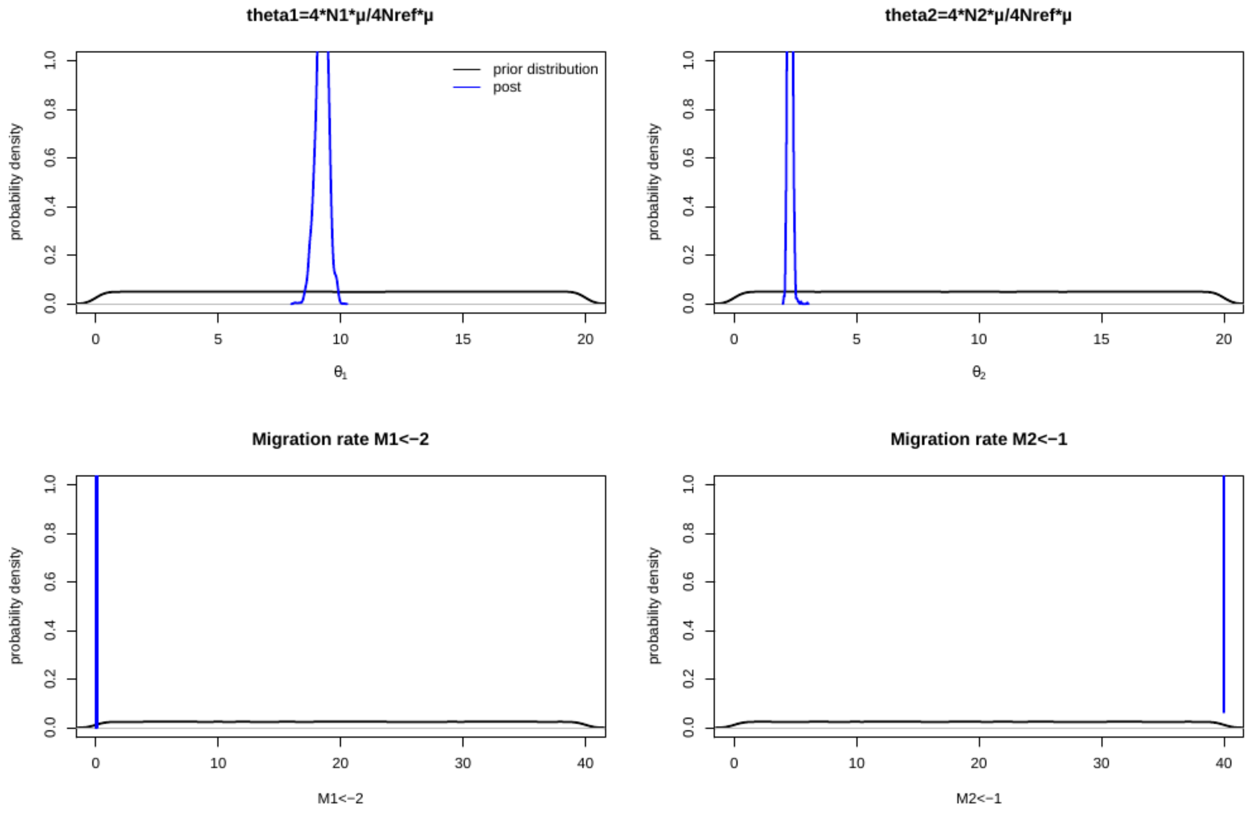

### B4DE – Best model = Equilibrium (Island model)

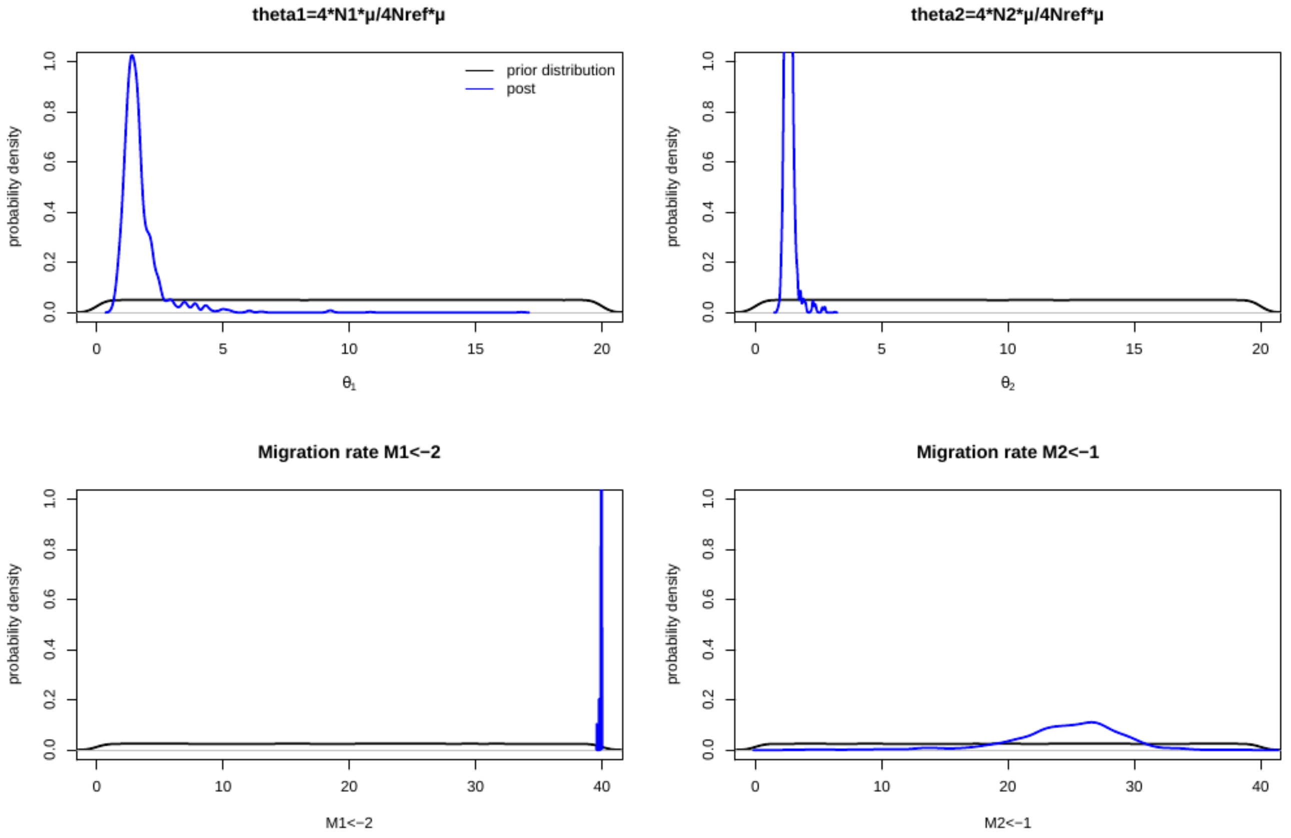

#### B5DE – Best model = Equilibrium (Island model)

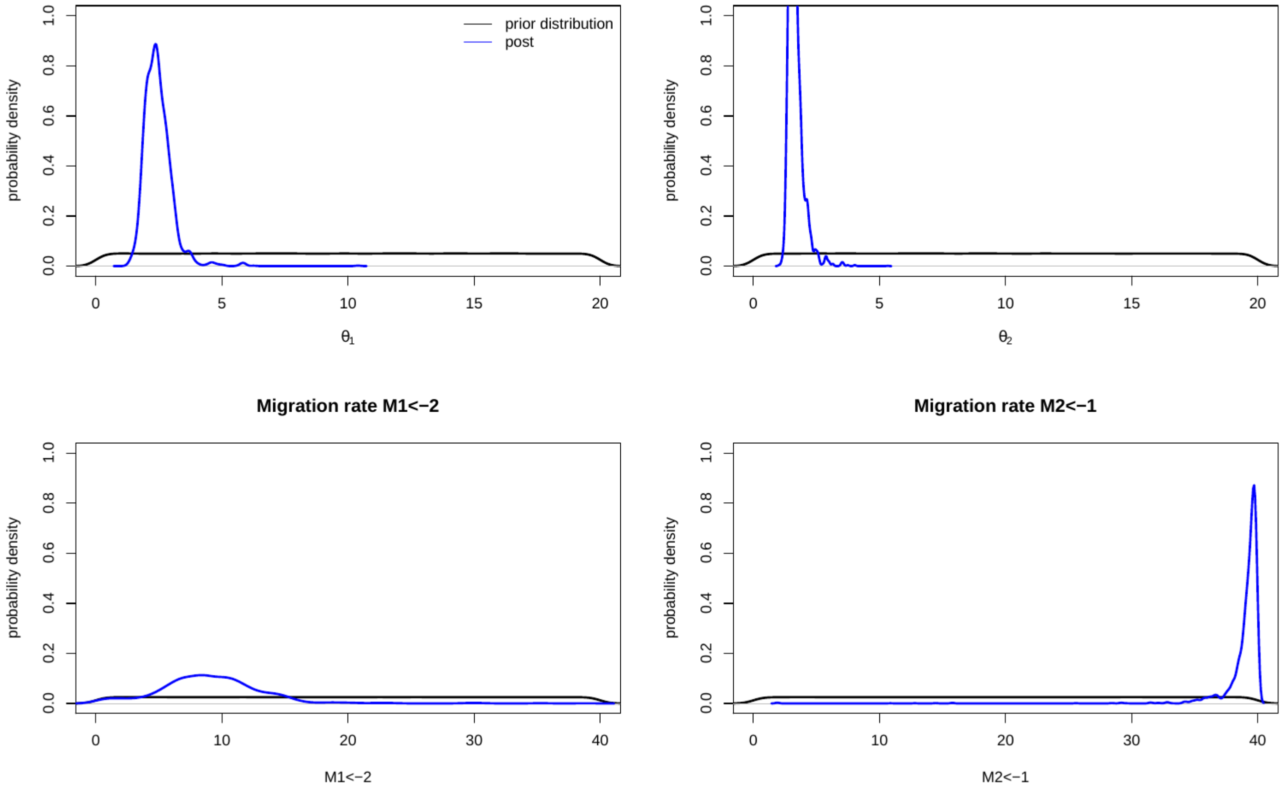


#### B6DE – Best model = Equilibrium (Island model)

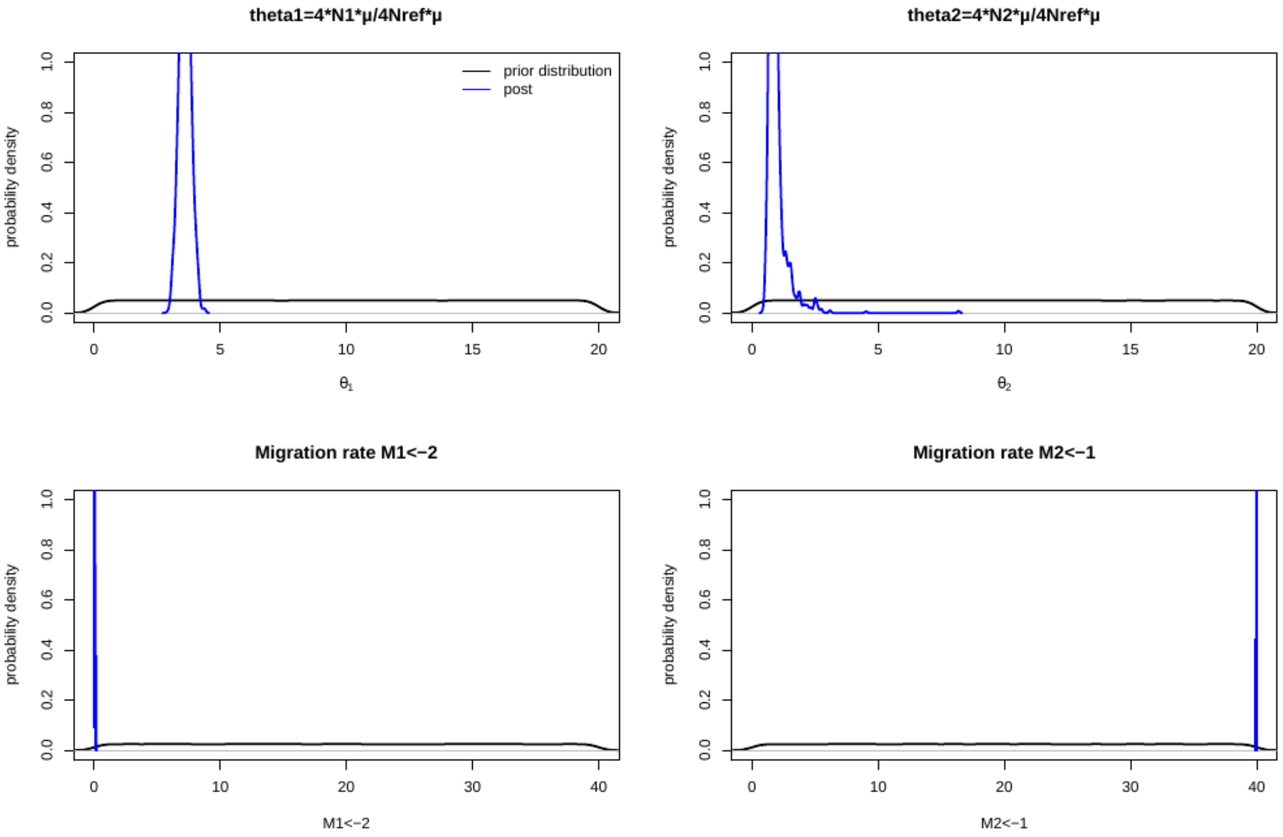
