## Supplementary Figure 3 for "Demographic history and genomics of local adaptation in blue tit populations"

**Supplementary Figure 3: Prior and posterior probabilities with larger bounds on migration rate.**

Prior and posterior probabilities of estimated parameters for each pair of population. Theta1 = 4N1/μ (scale mutation rate for the population 1), Theta2 = 4N2/μ (scaled mutation rate for population 2). ThetaA = 4Nanc/μ/. (scaled mutation rate for the ancestral population. All theta are scale by the theta of the reference population (4*Nref*μ). Migration rate $M_{ij}$ = 4N * ref *$m_{ij}$.

To reduce the exploration of the parameter space, we also reduce the lower and upper bounds on the scale effective population size (theta) based on our inferences in the first round of ABC analyses (Supplementary Figure 2).

#### A2DE – Second best model:  = Equilibrium (Island model)

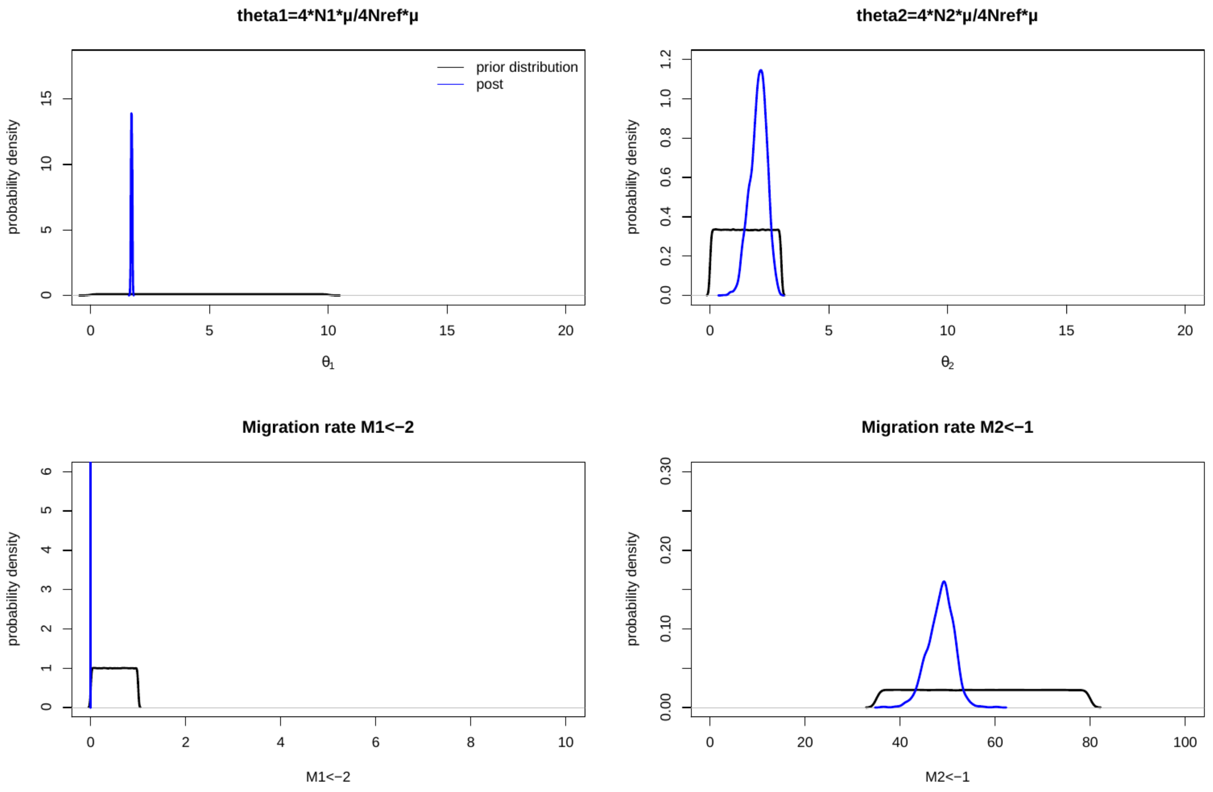

### B4DE – Best model = Equilibrium (Island model)

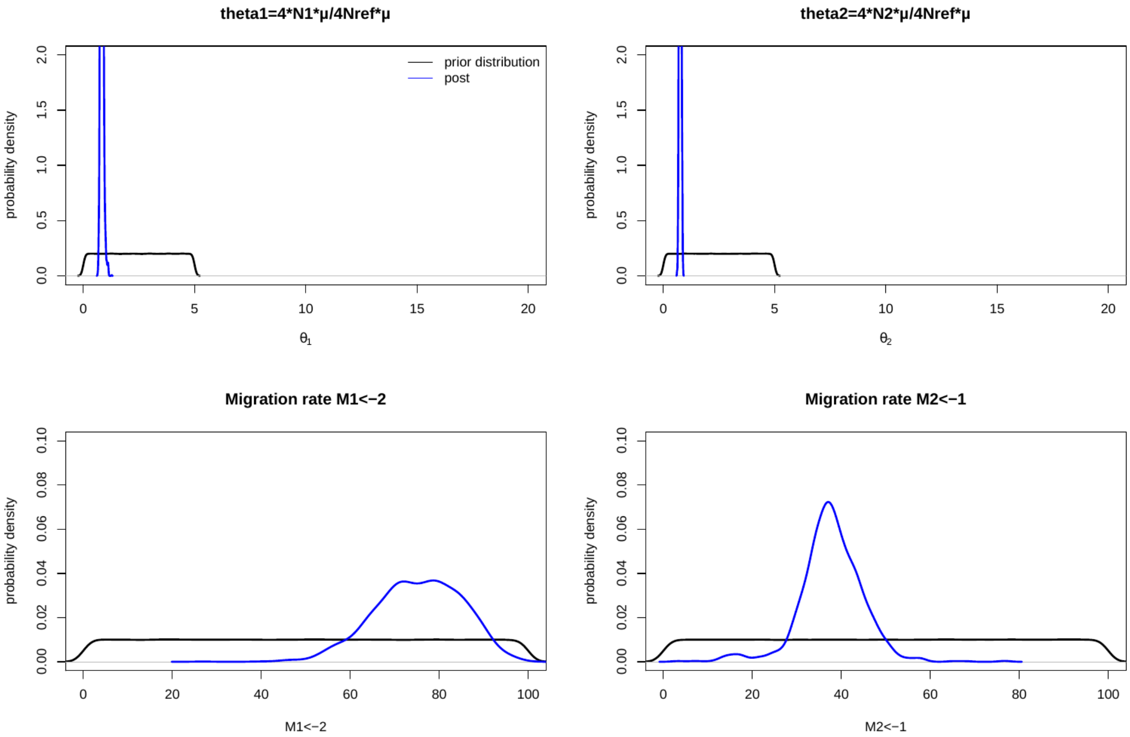

### B5DE – Best model = Equilibrium (Island model)

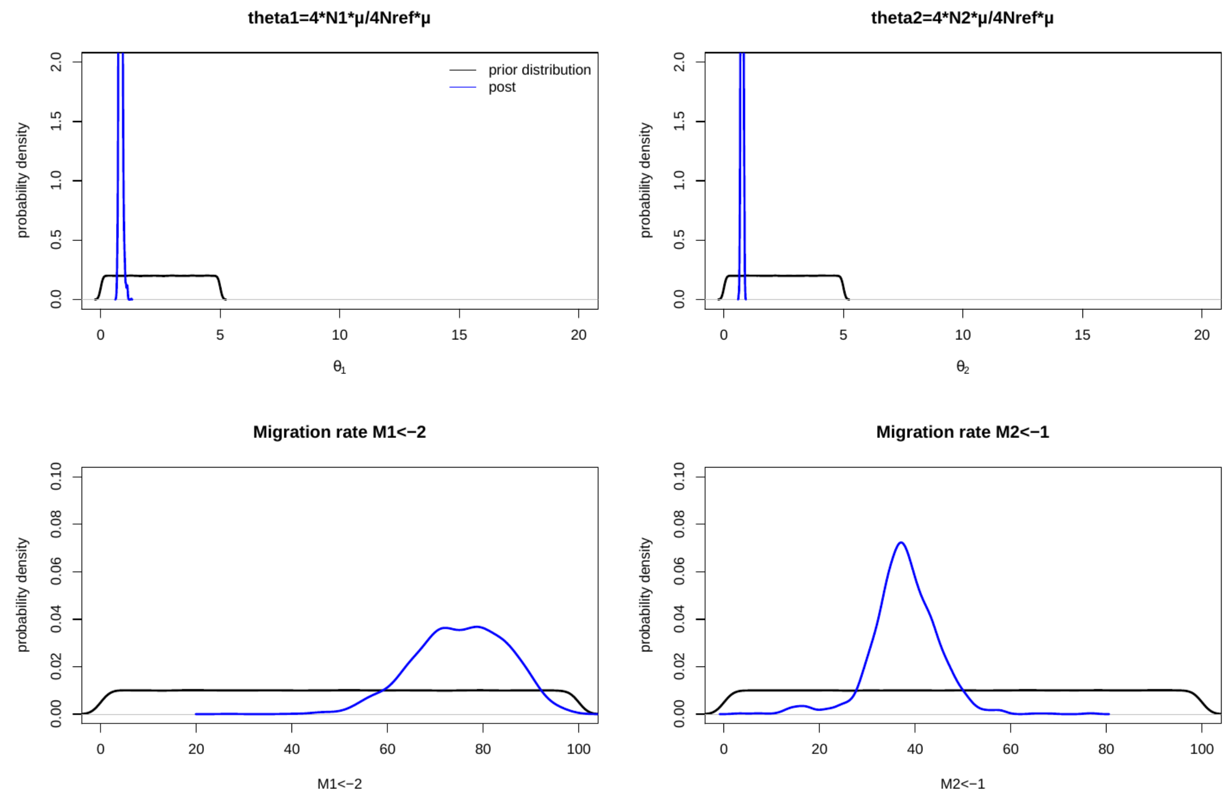


### B6DE – Best model = Equilibrium (Island model)

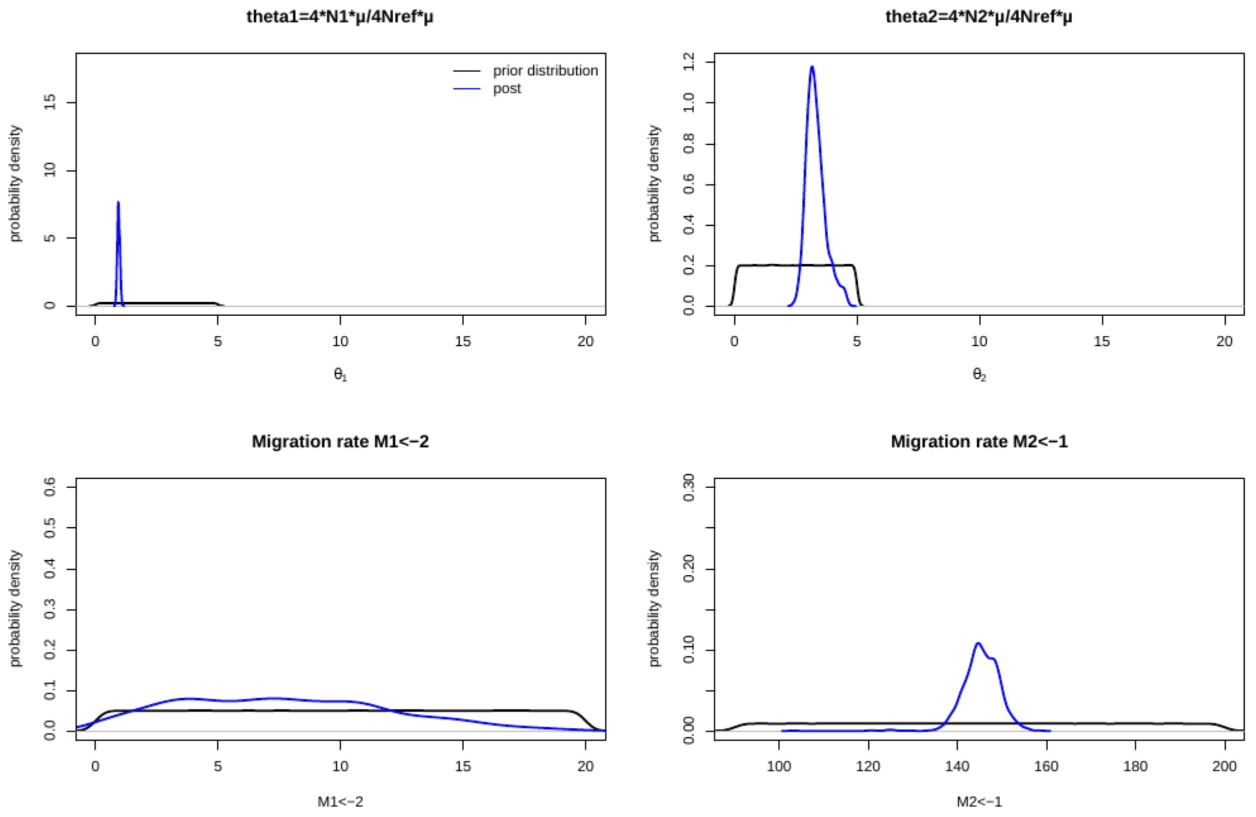
