## Supplementary Figure 4 for "Demographic history and genomics of local adaptation in blue tit populations"

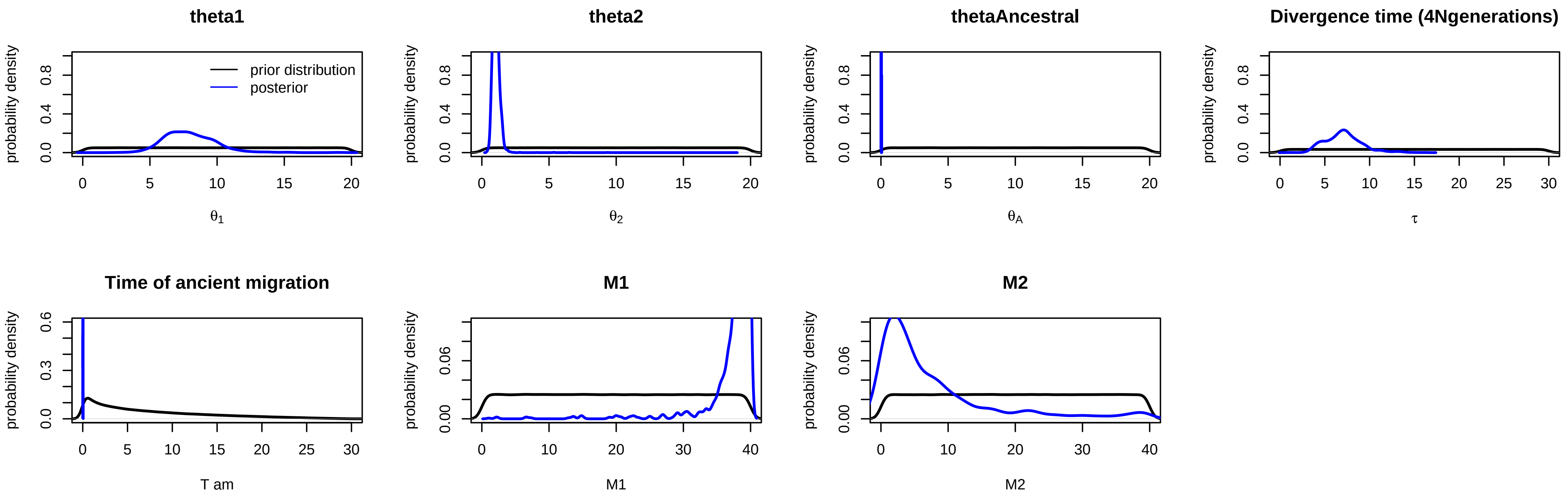


Supplementary Figure 4. Prior and posterior probabilities for models infering mainland-Corsica divergence
