## Supplementary Figure 5 for "Demographic history and genomics of local adaptation in blue tit populations"

Supplementary Figure 5. A) MAF histogram, B) LD decay with genomic distance (bp), and C) temporal Ne, for each populations.

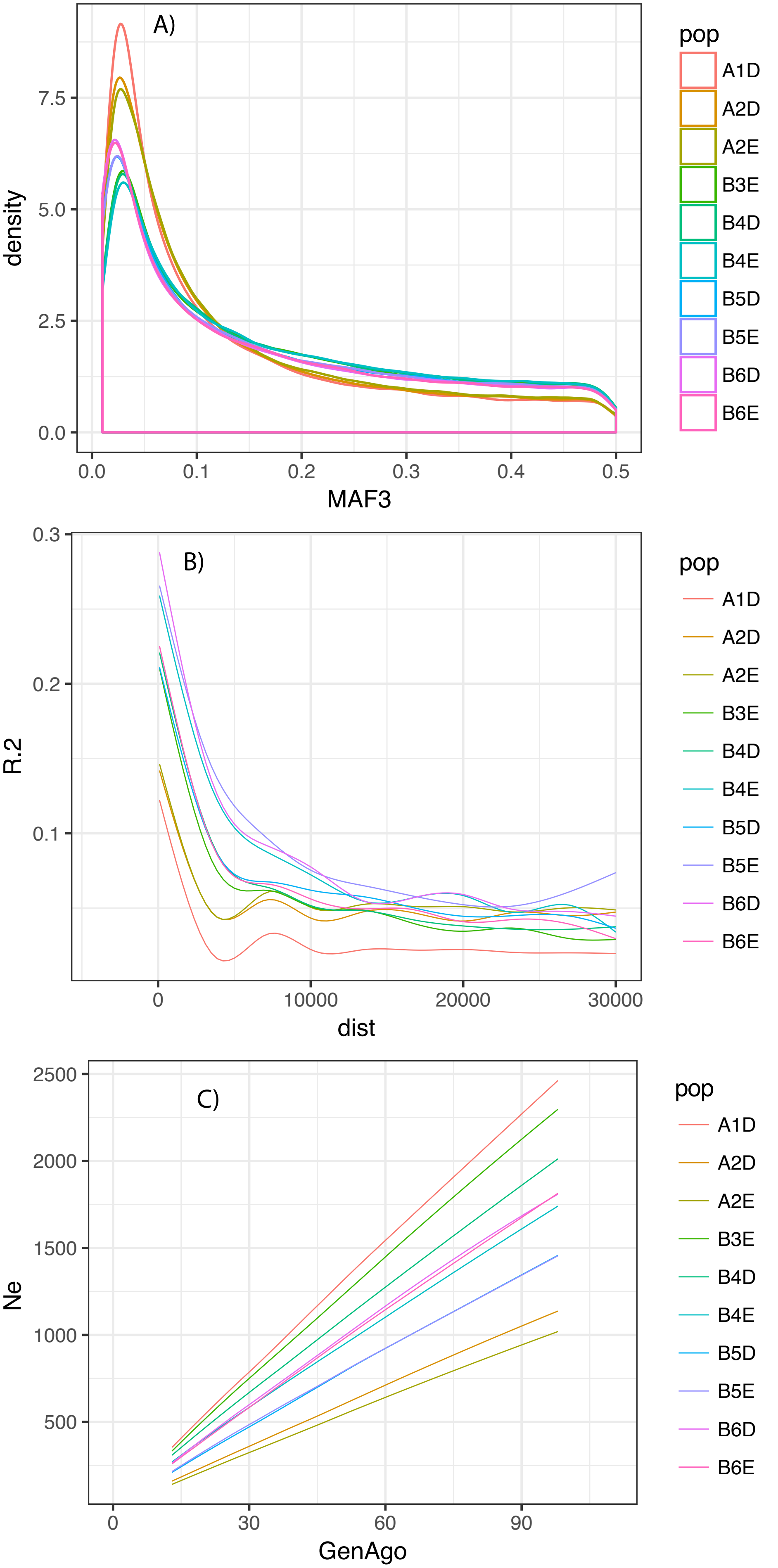
