## Supplementary Figure 6 for "Demographic history and genomics of local adaptation in blue tit populations"

Supplementary Figure 6. LD decay with genomic distance, for each populations, for the chromosomes 1, 2 and Z.

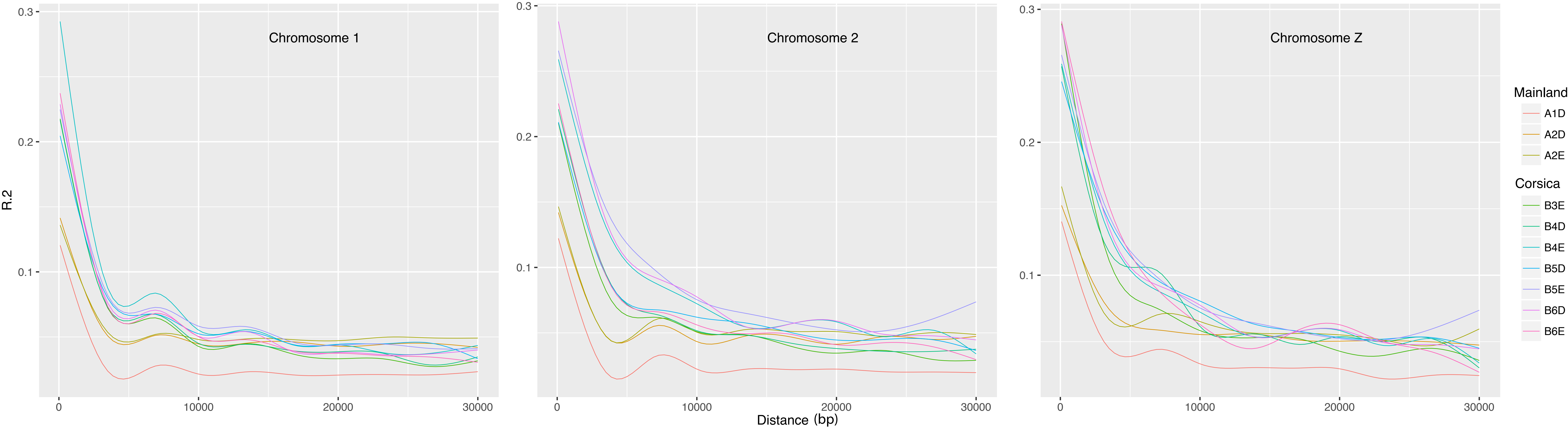
