## Supplementary Figure 9 for "Demographic history and genomics of local adaptation in blue tit populations"

Supplementary Figure 9. Fst as a function of Bayescan log10(BF) outputs for each SNP, for i) Corsica vs mainland and ii) evergreen vs deciduous populations.

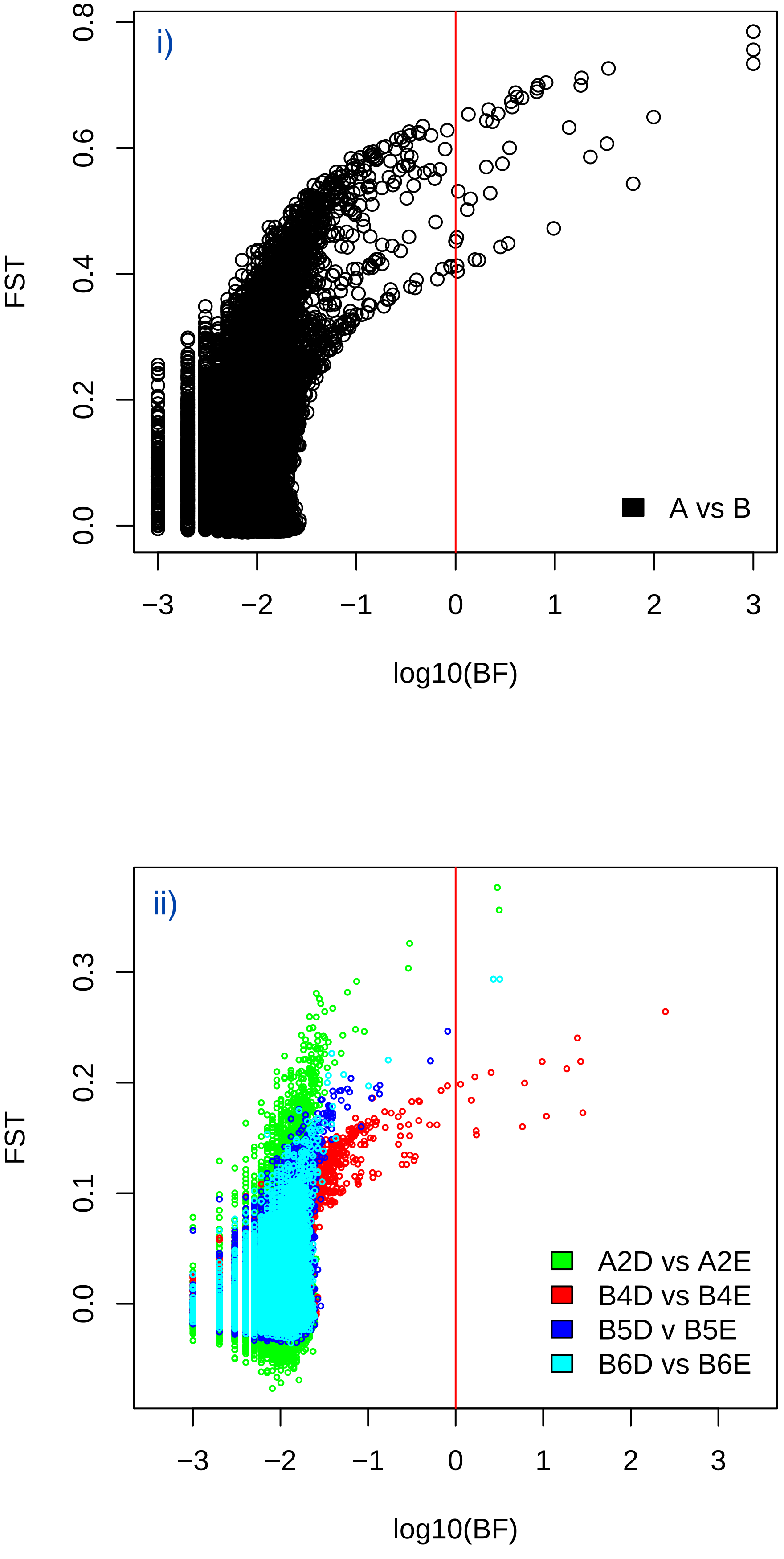
