## Supplementary Figure 10 for "Demographic history and genomics of local adaptation in blue tit populations"

Supplementary Figure 10. Histograms of SNP loadings for RDA i) constrained to test divergence between deciduous and evergreen populations and ii) constrained to test divergence between mainland and Corsica

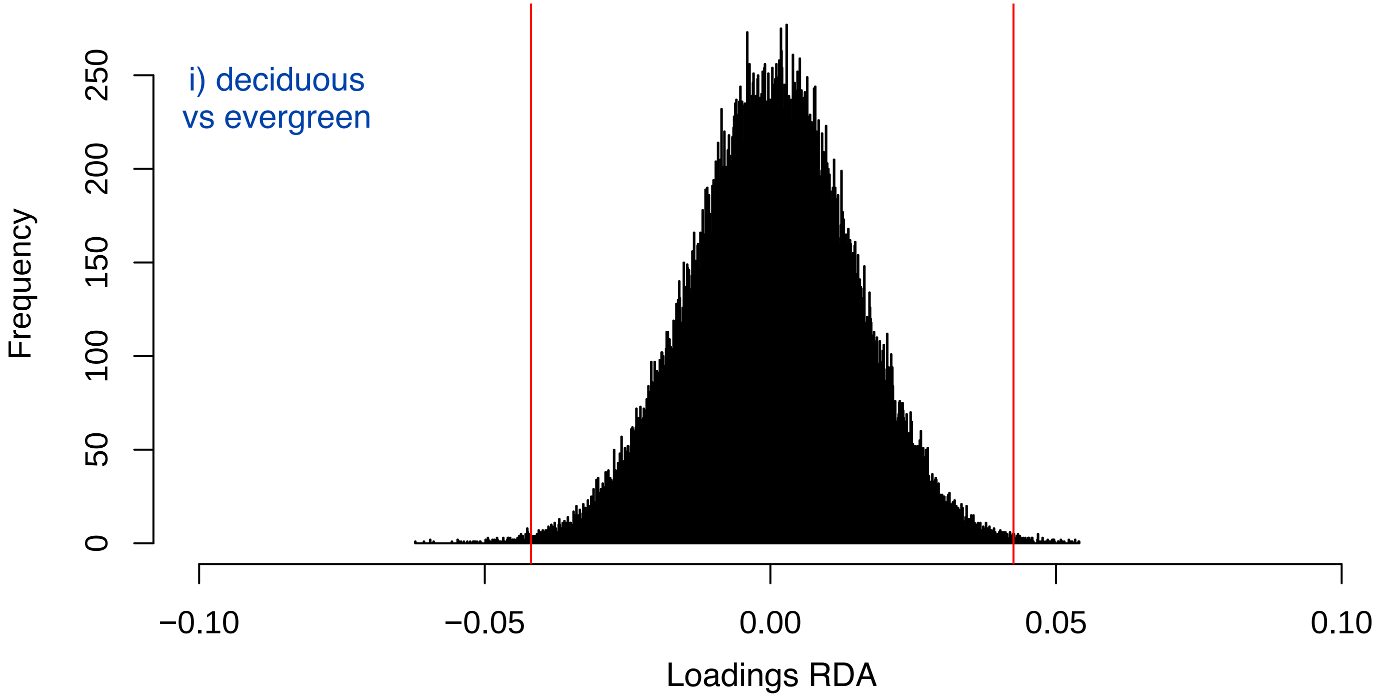


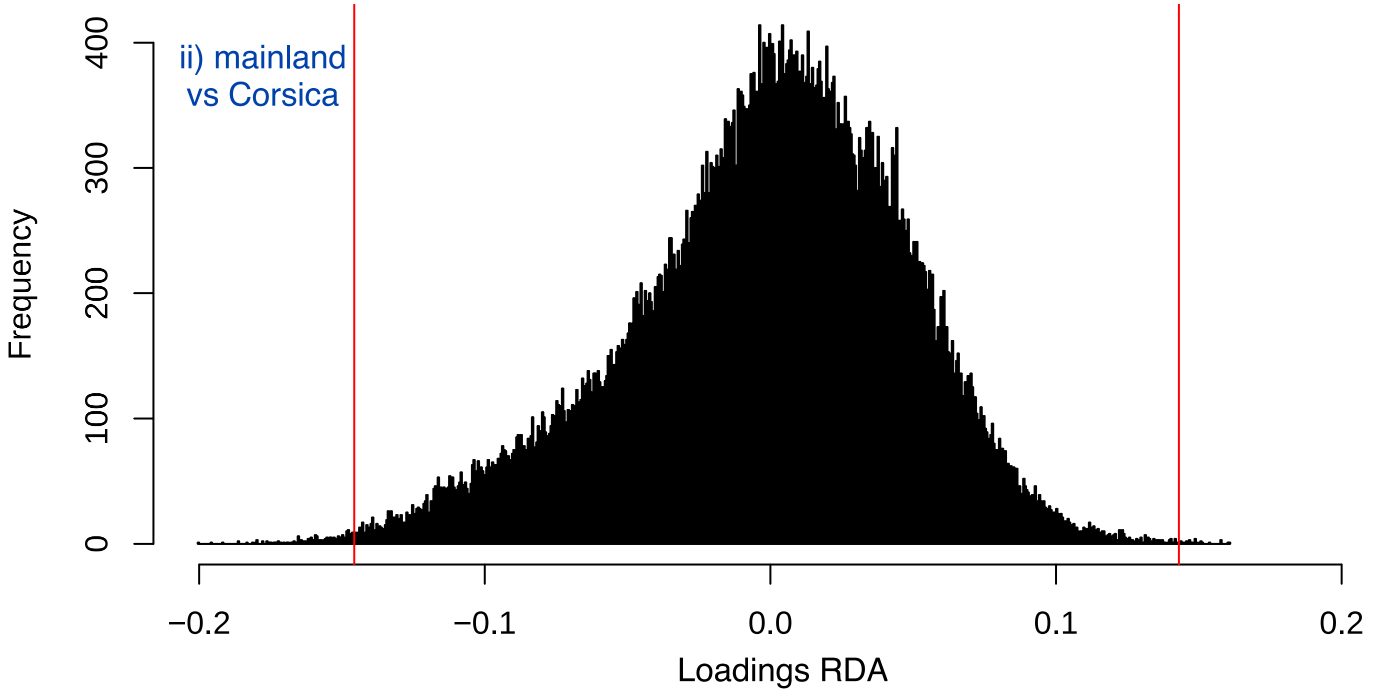
