## Supplementary Note 1 for "Demographic history and genomics of local adaptation in blue tit populations"

Six demographic scenarios were used for Approximate Bayesian Computation*:*

1) a model of panmixia (PAN),

2) a model of equilibrium corresponding to the island model with two populations (EQ),

3) a model of isolation with migration (IM),

4) a model of secondary contact (SC),

5) a model of divergence with migration during the first generations, *i.e.* ancestral migration (AM),

6) a model of strict isolation (SI).

*Prior and coalescent simulations:*

We used the coalescent simulator msnsam (Ross-Ibarra et al. 2008) a modified version of the coalescent simulator ms (Hudson 2002) under an infinite-site mutation model, and generated a total of 1x10^6^ coalescent simulations for each model with the same number of loci as those observed in each pair-wise dataset. We took advantage of haplotypic data from RADseq loci available from Stacks to simulate the data and used a series of custom scripts to filter those data to minimize missing rate and generate input files for coalescent simulations. For each pair of populations, we filtered for missing data < 20%, MAF > 0, read depth between 5-95% of the original distribution and removing the loci with a number of alleles < 95% of the distribution. The mutation rate was fixed to 1e^-8^ mutations/bp/generations.

In the absence of any prior information regarding parameters estimates, large and uninformative priors were used, as commonly done in the ABC literrature. We fixed Nref to 50 000 individuals and sampled uniformly *N_ANC_, N_Pop1_,* and *N_Pop2_*  on the interval [0-1,000,000]. Prior divergence time for IM, AM, SC and SI models were uniformly sampled on the interval [0-6,000,000] of generations, conditioning on *T*_am_ ,*T*_sc_ being sampled within this interval. The migration rate 4Nm was sampled on the interval [0-40] and independently in each population allowing for asymmetric gene flow. Priors were generated using a modified Python version of *priorgen* software (Ross-Ibarra et al. 2008) and a panel of 19 commonly used summary statistics (Fagundes et al. 2007; Roux et al. 2013) was computed using mscalc (Ross-Ibarra et al. 2008, 2009; Roux et al. 2011). These include the mean and standard variations of 1) the nucleotide diversity π (Tajima, 1989), 2) the net divergence between the two populations (DA), 3) the total divergence between populations (D_XY_) (Nei & Kumar, 2000), 4) the between population differentiation (F_ST_) value, 5) the number of fixed differences between populations (Sf); 6) the number of polymorphic sites private to each population (Sxpop1 and Sxpop2) and 7) the number of shared polymorphic sites (Ss); 8) and last Pearson’s R2 coefficient of correlation in π between the two populations. The whole pipeline is available at <https://github.com/QuentinRougemont/abc_inferences>**.**

*Model selection:*

Posterior probabilities of each model were obtained by comparing models using an ABC framewok. The best simulations (kept using a tolerance of 0.025% during the rejection step), were weighed by an Epanechnikov kernel peaking when *S*_obs_ = *S*_sims_. Posterior probabilities were then computed using fifty trained neural networks and 15 hidden networks in the regression using abc R package (Csillery et al. 2012). The feed-forward neural networks use nonlinear multivariate regressions and consider the model as an additional parameter to be inferred. The posterior probability was obtained after averaging over ten ABC replicates. Then, the robustness, defined as the probability P of correctly classifying a model M given a posterior probability threshold X, was assessed using pseudo-observed dataset (PODS) and computed using the following formula from (Roux et al. 2016). A total of 5,000 PODS from each model were used with parameters drawn from the same prior distribution as our previous coalescent simulation. To obtain the posterior probability of each PODS the ABC model choice procedure was run 5,000 time using the same parameter as our empirical data.

*Parameter estimation:*

Parameter estimation was obtained for the best model in each pairwise comparison after performing another set of 10,000,000 coalescent simulations with the exact same procedure as above. A logit transformation of the parameters and a tolerance of 0.001 was used to compute the posterior distribution. The neural network procedure with nonlinear regressions of the parameters on the summary statistics was used with 50 feed-forwards neural networks and 15 hidden layers, as implemented in the R pacakge ABC (Csillery et al. 2012).

Refinement of parameter estimates

In a number of pairwise comparisons, (i.e. A2D vs A2E; B4D vs B4E, B5D vs B5E, B6D vs B6E) we observed that the posterior distribution of estimated migration rate often reached the maximal upper bound of our initial prior. Therefore, we run another round of parameter estimation under the best model by (i) increasing the upper bound on the migration rate for all these models and (ii) fixing the theta values based on their posterior distribution inferred in the initial round of parameter estimations. This enabled us to refined our estimates of migration rates.
